# HUMAN SPINAL CORD ORGANOIDS REVEAL CELL INTERCALATION AS A CONSERVED MECHANISM FOR SECONDARY NEURULATION

**DOI:** 10.1101/2024.11.19.624296

**Authors:** José Blanco-Ameijeiras, Yara El Majzoub, Mar García-Valero, Mariana M Faustino, Elena Rebollo, Javier Macho-Rendón, Jorge Corbacho, Juan Ramón Martínez-Morales, Elisa Martí

**Author notes:** Corresponding author and **Lead contact** Elisa Martí. **INCLUSION AND DIVERSITY** We support inclusive, diverse, and equitable conduct of research.

## Abstract

Human pluripotent stem cells (hPSCs) have enabled major advances in neural organoid research, yet reconstructing spinal cord development in vitro remains challenging, as it requires mimicking the early environment of body axis elongation. Here, by exposing hPSCs to defined extrinsic signals, we directed their self-organizing capacity to generate organoids that recapitulate the transcriptional profile, cellular composition, and tissue architecture of the early human posterior spinal cord. Furthermore, we refined our culture system to more closely model the in vivo morphogenetic events of secondary neurulation. Using this approach, we identified cell intercalation—regulated by Yes-associated protein (YAP) activity—as a key morphogenetic mechanism driving de novo lumen formation and resolution. These biomimetic models provide a powerful platform to investigate the molecular and mechanical processes underlying human spinal cord development and offer new opportunities to elucidate the origins of neural tube defects, among the most common congenital defects.

## INTRODUCTION

The integration of stem cell biology and bioengineering has paved the way for the creation of in vitro organs. This field relies on pluripotent stem cells (PSCs), which can be directed into specific cell types through the careful regulation of signaling pathways. When combined with three-dimensional scaffolds, this method facilitates the formation of organoids—3D cellular structures that closely mimic the architecture of actual organs. Over the past decade, extensive research involving human PSCs has resulted in notable progress in the development of neural organoids. Groundbreaking studies have led to the creation of organoids that replicate neural structures such as the optic cup(Nakano et al., 2012), cerebral cortex(Kadoshima et al., 2013; Lancaster et al., 2017; Lancaster and Knoblich, 2014; Pasca et al., 2015), midbrain(Jo et al., 2016), hypothalamus(Kasai et al., 2020), and brainstem(Eura et al., 2020), all of which closely resemble their in vivo counterparts in terms of organization and function.

However, reconstructing spinal cord development in vitro has proven to be a complex challenge, as it demands a culture environment that closely resembles the early stages of body axis elongation and the associated signaling mechanisms. During embryonic development, the spinal cord originates from a population of bipotent stem cells termed neuro-mesodermal progenitors (NMPs)(Tzouanacou et al., 2009), which are organized within a transient embryonic structure known as the neural tube (NT)(Saade and Martí, 2025). Spinal cord morphogenesis differs along its anterior-posterior axis: most of the NT (including the regions that will develop into the brain and the anterior spinal cord) forms via primary neurulation, but the establishment of the posterior region of the spinal cord involves de novo formation in a process named secondary neurulation(Saade and Martí, 2025).

Human PSCs can be directed to form NMPs by mimicking key events of early embryogenesis in vitro(Gouti et al., 2017, 2014). These NMPs then organize themselves into spinal cord-like organoids, exhibiting several hallmark features of the spinal cord, such as the formation of patterned neuronal subsets with distinct dorsal-ventral identities, self-directed axial elongation, and gene expression profiles characteristic of specific spinal cord regions(Gribaudo et al., 2024; Libby et al., 2021; Xue et al., 2024). Additionally, these organoids can partially replicate NT morphogenesis through primary neurulation(Karzbrun et al., 2021; Lee et al., 2022; Rito et al., 2025). However, effective in vitro models of secondary neurulation, crucial for studying the origins of congenital NT defects (NTDs), are still lacking in the field.

Here we set to generate human spinal cord organoids, by exposing hPSCs to various extrinsic signals which play crucial roles in controlling the temporal and spatial regulation of gene expression necessary for building the NT along the anterior-to-posterior embryo axis(Simunovic et al., 2019; Turner et al., 2017). Our screening strategy demonstrated that the self-assembling properties of hPSC can be directed to generate organoids to mimic the developmental time window and the correct posterior axis identity of the early human embryo posterior spinal cord.

Moreover, to investigate the tissue dynamics of secondary neurulation, we refined our in vitro system to more closely mimic the in vivo morphogenetic process. In our posterior organoid model of secondary neurulation, we demonstrated that cell intercalation, regulated by the activity of Yes-associated protein (YAP), acts as a key morphogenetic mechanism for lumen formation. This indicates a conserved mechanism, similar to that observed in chick embryos(Gonzalez-Gobartt et al., 2021). Collectively, these findings establish the first human organoids with a posterior spinal cord transcriptional identity and provide a reliable model of secondary neurulation, which may advance our understanding of the biological basis of NTDs.

## RESULTS

### Generation of human neural organoids with different anterior-posterior identities

During neural tube (NT) formation and elongation, progenitor cells are sequentially exposed to a variety of extrinsic signals, including members of the TGFβ/BMP, WNT, FGF and RA signalling families (Figure 1A). These signals play crucial roles in controlling the temporal and spatial regulation of gene expression, which in turn regulates lineage restriction and the generation of specific cell identities necessary for building a functional nervous system(Olivera-Martinez et al., 2012; Ribes et al., 2009; Wilson and Hemmati-Brivanlou, 1995; Wilson et al., 2009). Here we have set to mimic the embryo environment to guide human pluripotent stem cells to build neural tube (NT) organoids.

**Figure 1:**
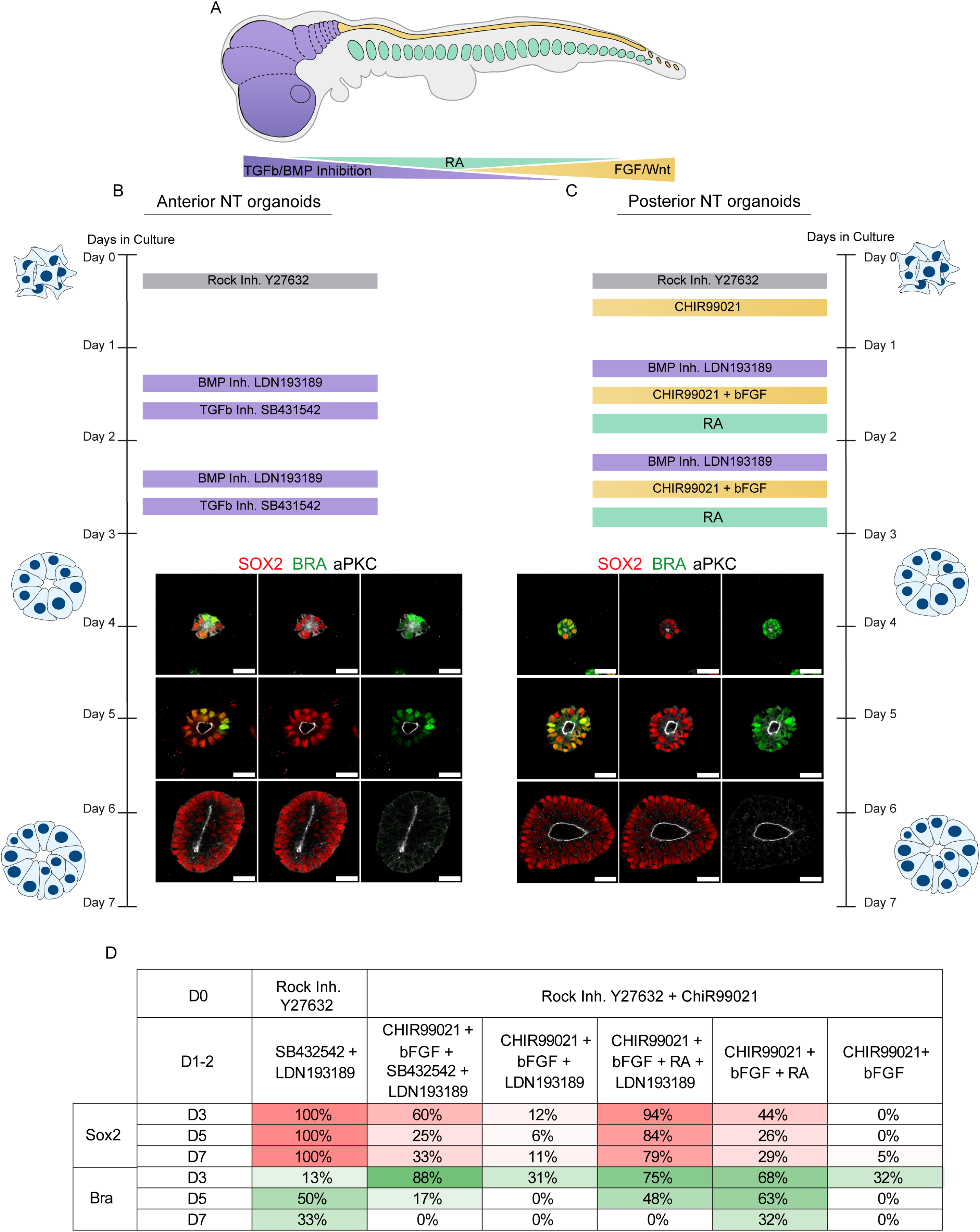
Generation of neural organoids with different anterior-posterior identities (see also Figures S1-2). **(A)** Schematic representation of an embryo; the forming brain is shown in purple, the spinal cord in yellow and the somites are shown in green. Gradients of morphogenetic signals are shown. **(B, C)** Schematic representation of organoid culture, guiding drugs are indicated at the appropriate days in culture. Selected images of anterior (B) and posterior (C) organoids stained for SOX2 (red), BRA (green), aPKC (grey). Scale bars = 30 µm. **(D)** Quantification of organoids expressing the indicated markerş SOX2 (red) and BRA (green), after 3, 5 and 7 days (D) in culture. Guiding drugs are indicated in each column, as well as days (D) of treatments.

Members of the TGFβ/BMP signalling pathway are known to be present and active during neural induction and neural plate formation(Wilson and Hemmati-Brivanlou, 1995). *In vitro* studies conducted in both 2D(Chambers et al., 2009; Verrier et al., 2018) and 3D cultures(Haremaki et al., 2019; Zheng et al., 2019) have reported that double inhibition of TGFβ and BMP leads to rapid neural conversion. We initially seeded human embryonic stem cells (hESC; RUES2) on matrigel and cultured them in neural induction medium (N2B27) with transient inhibition of BMP and TGFβ signalling (48-hour treatment with the BMP type I receptor ALK1-3 inhibitor LDN193189, and the TGFβ type I receptor ALK4,5,7 inhibitor SB431542). This resulted in the organization of multicellular cysts, termed neural organoids, consistently composed of SOX2+ cells (Figure 1B, D). Concomitantly with the neural lineage restriction of hESCs, these SOX2+ cells underwent epithelialization, as evidenced by the apical localization of aPKC, and self-organized around a single central lumen (Figure 1D). A 48-hour pulse of TGFβ/BMP inhibition mimicked neural induction, as reported(Ogura et al., 2018; Simunovic et al., 2019), but some neural organoids still expressed Bra, likely because the inhibition period was shorter than in earlier studies (Figure 1B, D).

Our aim was to establish a combination of extrinsic signals that closely mimics the environment of the elongating embryo, and hence the posterior spinal cord formation (Figure 1A). This combination should effectively guide the conversion of NMP cells, as monitored by the dynamic expression of SOX2/BRA, and consecutively promote the mesenchymal-to-epithelial transition and the self-organization around a single lumen. To begin to mimic the signalling environment of the posterior embryo, we exposed the hESC to the WNT agonist CHIR (CHIR99021) for initial 24 hours followed by a 48 hours pulse along with basic FGF (bFGF)(Edri et al., 2019; Takemoto et al., 2006). Consistent with previous studies in mouse ES cells and mouse epiblast cells, this resulted in the transient presence of BRA+ cells(Edri et al., 2019; Koch et al., 2017). Moreover, this condition appeared to be insufficient to guide neural progenitor conversion, reflected by the failure of the maintained SOX2+ expressing cells, the lack of complete a MET, and the failure to organize a central lumen (Figure 1D; Supplementary Figure 1F). In vivo, the neural specification from NMPs is regulated by Noggin (a BMP antagonist) secreted by the notochord(McMahon et al., 1998). To mimic this embryonic environment, we next exposed hESCs to WNT/bFGF along with TGFβ and/or BMP inhibition, to burst neural induction. Results showed that, in the presence of WNT/bFGF, neither BMP inhibition alone, nor the double TGFβ/BMP inhibition, were sufficient to promote a robust neural conversion, since the percentage of organoids fully formed by SOX2+ cells were low in both culture conditions (Figure 1D; Supplementary Figure 1B,C).

In the elongating embryo, Retinoic Acid (RA) mainly secreted from the somites (Figure 1A) plays an instructive role by opposing WNT activity and regulating the balance between the NMP cell fate and neural identity(Del Corral et al., 2003). Hence, we next exposed hESCs to neural induction medium supplemented with WNT/bFGF, BMP inhibition, and RA for 48 hours. After three days in culture, hESCs efficiently formed small organoids consisting of NMP-resembling cells expressing SOX2/BRA (Figure 1 C,D). The temporal evolution of these NT organoids also resembled the lineage restriction of NMP cells *in vivo*, with the presence of BRA+ cells gradually decreasing. Furthermore, these organoids efficiently self-organized to form a single central lumen (Figure 1 C,D). We next tested whether, in the presence of RA, BMP inhibition was dispensable for the formation of these neural organoids. After three days of growing hESCs in medium supplemented with WNT/bFGF and RA for 48 hours, organoids were formed where the presence of BRA+ cells decreased along the time in culture, while that of SOX2+ cells was low (Figure 1 D; Supplementary Figure 1 E), altogether indicating the relevance of BMP inhibition for an efficient neural induction.

Next, to investigate whether the caudal limit observed in mouse ESC differentiation without WNT signalling is conserved in human cells, we examined organoids generated from hESCs cultured under conditions that inhibited TGFβ/BMP activity. In these organoids, the presence of CDX2+ cells was rare, indicating an anterior identity of these organoids (Supplementary Figure 1A). However, when hESCs were cultured in the presence of WNT/bFGF, BMP inhibition, and RA, organoids consistently and robustly exhibited the presence of CDX2+ cells, indicative of a posterior (spinal cord) identity of these organoids (Supplementary Figure 1D). In summary, we generated neural organoids exhibiting a posterior spinal cord identity using an in vitro protocol that mimics the elongating embryonic environment and that, compared with previous methods, requires a shorter treatment pulse to establish this identity(Iyer et al., 2022; Maury et al., 2015; Tang et al., 2022).

Finally, to demonstrate the robustness of our protocol in generating neural organoids, we utilized induced pluripotent stem cell (human iPSC) lines, namely Kolf2-C1 and CRTD1. We cultured these iPSCs in neural induction medium with different conditions to guide the generation of organoids with distinct anterior and posterior identities. When iPSCs were cultured with transient inhibition of TGFβ/BMP signalling for 48h, they were directed to form NT organoids with anterior neural identity. In contrast, iPSCS cultured in neural induction medium with WNT/bFGF, BMP inhibition, and RA formed organoids where the temporal evolution resembled that of lineage restriction of NMP cells in vivo (Supplementary Figure 2A-G). These findings demonstrate that the self-assembling properties of human pluripotent stem cells can be directed to generate NT organoids with defined anterior and posterior identities.

### Transcriptome profiling of neural organoids revealed different anterior and posterior identities

To better define the identity of the *in vitro*-generated neural organoids, we performed bulk RNA sequencing (RNA-seq) using three biological replicates of anterior (A) and three replicates of posterior (P) neural organoids (Figure 2A). These organoids were generated from human embryonic stem cells (hESC; RUES2) and were grown *in vitro* for 6 days. During this culture period, the organoids acquired the A and P identities, respectively, but neurogenesis was not yet initiated.

**Figure 2:**
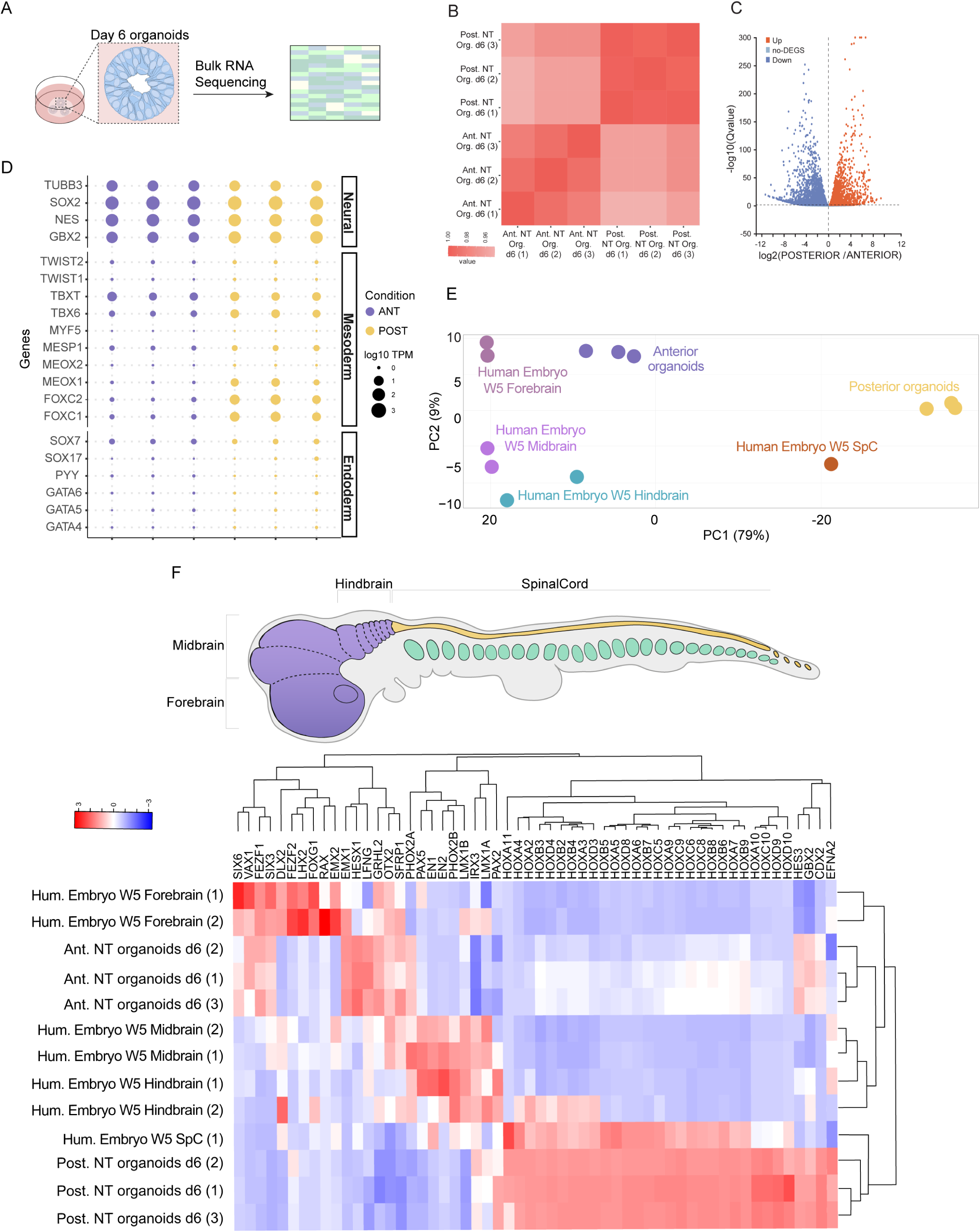
Anterior and Posterior neural organoids show different transcriptome identity. **(A)** Schematic representation of organoid culture and RNA extraction for bulk RNA Sequencing. **(B)** Heatmap of the pairwise Pearson correlation coefficients of gene expression in the three replicates for both Anterior and Posterior organoids. **(C)** Volcano plot representation of differential expression analysis of genes in Posterior and Anterior NT organoids. **(D)** Bubble plots the expression levels of selected neural, mesodermal and endodermal genes in the anterior (blue) and posterior (yellow) organoid samples. **(E)** Principal Component Analysis (PCA) of RNAseq-based expression data of selected genes for forebrain (purple), midbrain (lilac), hindbrain (purple) and spinal cord (mustard) samples isolated from human embryos (week 5 of development) and anterior (blue) and posterior (yellow) neural organoid samples. **(F)** Schematic representation of an embryo; the Anterior forming brain is shown in purple, Posterior forming spinal cord in yellow, and the somites are shown in green. **(G)** Heatmap of the expression of the AP domain-associated selected genes in human forebrain, midbrain, hindbrain and spinal cord samples isolated from human embryos (week 5 of development) and compared to the Anterior and Posterior neural organoid samples. In the x-axis it is represented the hierarchical clustering of the AP domain-associated genes. In the y-axis it is represented the hierarchical clustering of the samples.

We employed the DNASEQ platform from BGI to identify 17,462 genes, with an average mapping ratio to the reference genome of 98.03% (Reference Genome Version: GCF_000001405.39_GRCh38.p13). The Pearson correlation coefficients between samples revealed low gene expression variability among the three A and the three P organoid samples (Figure 2B). A total of 8,728 differentially expressed genes (DEGs) were identified when comparing the transcriptomes of A and P organoids, confirming the distinct identities of these two types of neural organoids (Figure 2C).

We first examined the overall tissue identity of these organoids by assessing the combinatorial expression of a curated set of marker genes previously used to classify tissue types in early human embryos(Rayon et al., 2021; Zeng et al., 2023) (Figure 2D). This analysis revealed that the generated organoids were primarily composed of neural cells. A mesodermal component was detected in the posterior organoids, consistent with the neuro-mesodermal identity of early spinal cord progenitors. In contrast, endodermal gene expression was largely absent from both anterior and posterior neural organoids.

Next, to investigate the antero-posterior identity of the neural organoids, we mapped anterior (A) and posterior (P) samples onto the spatial transcriptome of neural tube cells from early human embryos at Carnegie stage 7 (week 3 of development), which encompasses the forebrain, midbrain, hindbrain, and spinal cord(Zeng et al., 2023). In A organoid samples, we observed strong expression of *SIX3* and *OTX2*, consistent with a forebrain-like identity, whereas these genes were nearly absent in P samples (Supplementary Fig. 3A). Midbrain-associated genes, including *EN1* and *FGF17*, were expressed at low levels in both A and P organoids. By contrast, hindbrain- and spinal cord–defining genes were expressed at higher levels in P than in A organoids, consistent with their posterior identity (Supplementary Fig. 3A). Moreover, HOX gene expression, which reflects the established antero-posterior patterning of the hindbrain and spinal cord, was largely restricted to posterior organoids (Supplementary Fig. 3A).

To refine the spatial characterization of A and P organoids, we next compared our samples with RNA-sequencing data from forebrain, midbrain, hindbrain, and spinal cord samples of human embryos at Carnegie stage 14 (corresponding to week 5 of development), a stage comparable to mouse embryonic day 10, when neurogenesis in the spinal cord is just beginning(Lindsay et al., 2016). Here, we examined the expression levels of a diagnostic panel of 41 genes that are specifically expressed along the antero-posterior axis of the developing CNS in distinct domains(Akin and Nazarali, 2005; Basson et al., 2008; Deng et al., 2011; de Vries et al., 2020; Hirata et al., 2001; Kumamoto and Hanashima, 2017; Porter et al., 1997; Yan et al., 2011). Principal components analysis (Figure 2E) and hierarchical clustering (Figure 2F) of expression profile for all samples confirmed that A organoid samples group together with the anterior nervous system samples, including CS14 forebrain, midbrain, and hindbrain controls, whereas P organoid samples were closer to the CS14 spinal cord control.

Detail examination of individual markers revealed that although anterior organoids exhibited similarity with forebrain samples, genes expressed in the dorsal telencephalon and neural retina domains, such as *EMX2*, *RAX2*, *FOXG1*, *LHX2*, *FEZF2*, and *DLX2* show low expression levels in the organoids (Figure 2F). This finding suggests that our culturing conditions did not favour the *in vitro* specification of the precursors towards dorsal retina and forebrain identities. Instead, they directed them towards ventral midbrain and retinal (i.e., optic stalk) derivatives, as indicated by the expression of genes like *VAX1*, *SIX6*, *SIX3*, or *FEZF1* (Figure 2F).

On the contrary, the gene expression clustering of P organoids samples revealed the remarkable absence of brain-specific gene markers, while these neural progenitors are capable of generating a full range of anterior-posterior spinal cord derivatives (Figure 2F). As such, the P organoids expressed both anterior (e.g., *HOX2-4*) and posterior *HOX* (e.g., *HOX9-10*) spinal cord genes, as well as very distal markers such as *CDX2* or *EFNA2*.

Next, we assessed the dorso-ventral (DV) identity of the posterior NT organoids by examining a diagnostic panel of 35 DV markers(Alaynick et al., 2011; Rayon et al., 2021) (Supplementary Figure 3B, C). This analysis revealed that in contrast to the CS14 spinal cord controls, which expressed a full range of DV markers, our posterior NT organoids have a dorsal identity, with most ventral genes (e.g. *SHH*, *NKX2.2*, or *OLIG2*) being almost absent (Supplementary Figure 3 B, C). However, this dorsal identity of the organoids reflects an early developmental stage in spinal cord formation, and it is consistent with our culture condition which is lacking any ventralization signal such as Sonic hedgehog(Saade and Martí, 2025). Hence, for these early organoids to form a bona fide spinal cord biomodel, symmetry breaking is further required by the addition of ventralizing signals such as Sonic hedgehog to the culture media, as previously reported(Krammer et al., 2024).

Importantly though, to our knowledge these are the first neural organoids exhibiting a robust spinal cord transcriptome identity, including the most posterior spinal cord, at the regions were the NT is developed through the morphogenetic process called secondary neurulation (SN)(Saade and Martí, 2025).

### Neural organoids are organized by cells exhibiting cell architecture and dynamics resembling those of the early neural tube progenitor cells

Regardless of the differential origin along the anterior to posterior axis of the NT, the architecture of the early embryonic NT (midbrain/hindbrain and spinal cord) is conserved along its entire A P axis, consisting on a hollow tube built by highly polarized neural progenitor cells (NPCs). We next set to evaluate whether the cellular and subcellular organization of A and P neural organoids, mimic the embryonic NT cell architecture and dynamics.

During NT formation in the chick and mouse embryo NPCs elongate(Saade and Martí, 2025). Hence, we first compared the cell shape of hESCs cultured in the presence of WNT/bFGF (resulting in a mesenchyme-like tissue organization), to those cultured in neural induction medium guided for either P or A identities. The results demonstrated a significant elongation of NPCs, in both neural organoids. The major axis/minor axis ratio of the cell shape was increased in NPCs of neural organoids (mesenchymal organoids median ± IQR major axis/minor axis = 1.48 ± 0.48; P organoids median ± IQR major axis/minor axis = 3.88 ± 1.97; A organoids median ± IQR major axis/minor axis = 4.37 ± 1.71; Figure 3 A, B and Supplementary Figure 4A, B)

**Figure 3:**
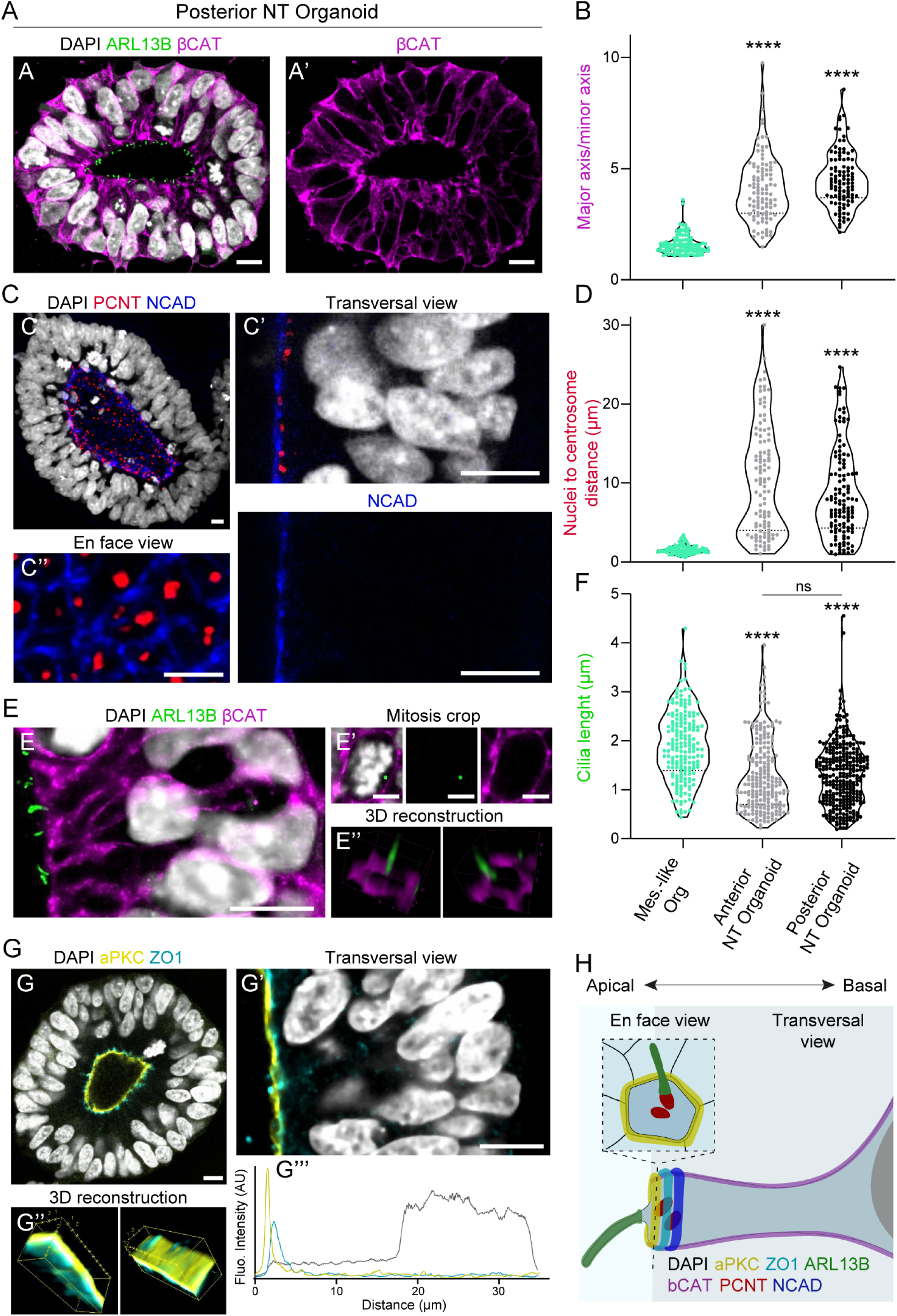
Posterior neural organoids are composed by epithelial polarized cells. **(A)** Selected images of Posterior (day 7) organoids stained for the ciliary membrane marker ARL13B (green) and the junctional protein βCatenin (βCAT, magenta), DAPI (white). Selected images of the βCAT staining alone are shown in **A’**. Scale bars = 10 µm. **(B)** Plots major axis/ minor axis of the cell (aspect ratio) in organoids with mesenchymal-like identity, Anterior, and Posterior organoids (horizontal bold lines show the median; n=100, 100 cells from 5 organoids; ****p<0.0001 Kruskal-Wallis test and Dunn’s multiple comparison test). **(C)** Selected images of Posterior organoids stained for the centrosome marker pericentrin (PCNT, red) and the junctional complexes protein N-Cadherin (NCAD, blue). Higher magnifications of transversal views are shown in **B’**. Scale bars = 10 µm. *En face* views is shown in **B’’**. Scale bars = 5 µm. **(D)** Plots nuclei to centrosome distance in cells from organoids with mesenchymal identity, Anterior, and Posterior organoids (horizontal bold lines show the median; n=187 cells from 5 organoids; ****p<0.0001 Kruskal-Wallis test and Dunn’s multiple comparison test). **(E)** Selected images of transversal views of Posterior organoids stained for ARL13B (green) and βCAT (magenta), DAPI (white). Scale bars = 10 µm. Higher magnifications of mitosis in transversal views are shown in **E’**. Scale bars = 5 µm. 3D reconstructions of the apical end foot of the cells are shown in **E’’**. **(F)** Plots cilia length in cells from organoids with mesenchymal identity, Anterior, and Posterior organoids (horizontal bold lines show the median; n=100, 100 cells from 5 organoids; ****p<0.0001 Kruskal-Wallis test and Dunn’s multiple comparison test). **(G)** Selected images of transversal views of Posterior organoids stained for the fate-determining factor aPKC (yellow) and the zonular protein ZO1 (cyan). Higher magnifications of transversal views are shown in **G’**. Scale bars = 10 µm. 3D reconstructions of the apical surface of the epithelium are shown in **G’’**. (**G’’’**) Plots show the fluorescence intensity to the distance from the organoid lumen (zero). Yellow line represents aPKC, cyan line represents ZO1 and grey line represents DAPI (white). **(H)** Scheme of polarized neuroepithelial cell in transversal and *en face* views. The centrosome (red) is apically localized. Cilia (green) are nucleated from the centrosomes pointing to the lumen of the tissue (red dots). Apical polarity components are organized in the apical end foot, being aPKC (yellow) apical to ZO1 (cyan) and NCAD (blue). βCAT (magenta) is localized in the cell membrane.

During NPCs elongation, the perinuclear centrosome relocates apically, as indicated by its alignment with N-Cadherin expression along the NT lumen. *En face* imaging of organoids revealed the localization of N-Cadherin to the NPC apical belt and the central localization of Pericentrin (PCNT+) labelled centrosomes (Figure 3 C, D). Quantification of the distance between the PCNT+ labelled centrosomes and the nucleus showed their apical localization in elongated NPCs of both in both A and P organoids (median ±IQR distance mesenchymal cells = 1.46 ±0.63 μm vs P organoids NPCs = 7.21 ±7.55 μm, and A organoids NPCs = 10.18 ±11.54 μm; Figure 3 C,D and Supplementary Figure 4 C, D). The distance of PCNT+ centrosomes from the nucleus exhibited wide variation, which could be attributed to the onset of interkinetic nuclear migration (INM), a process that separates or brings together the centrosome and the nucleus depending on the phase of the cell cycle, similar to what occurs in the developing NT *in vivo*. These findings indicate that our organoids display cellular and subcellular organization resembling that of the embryonic NT, including the elongation of NPCs and the apical localization of centrosomes in relation to the nucleus(Saade et al., 2018).

NPCs are characterized by the presence of a single primary cilium at their apical surface, which is nucleated by the basal body(Blanco-Ameijeiras et al., 2022), and extends into the lumen of the NT in the developing embryo. In both P and A organoids, the lumen surface was found to be decorated with primary cilia, identified by the expression of the protein ADP-ribosylation factor-like 13b (ARL13B), which specifically associates with the ciliary membrane(Paridaen et al., 2013; Saade et al., 2017) (Figure 3 E). The length and dynamics of these primary cilia in human NPCs generated *in vitro* were similar to those observed in the chick embryo NT(Saade et al., 2020). The length of Arl13b-labeled cilia protruding from the centrosome at the lumen of organoids was measured and showed comparable values (median ± IQR length: mesenchymal cells = 1.86 ± 0.98 μm; P organoids NPCs = 1.29 ± 0.92 μm; A organoids NPCs = 1.11 ± 1.03 μm; Figure 3 E, F and Supplementary Figure 4 B). During mitosis in organoids, ARL13B was observed to accumulate near one of the centrosomes that nucleated the mitotic spindle (Figure 3 E’). This finding suggests that, similar to observations in the developing mouse neocortex(Paridaen et al., 2013) and the chick embryo NT(Saade et al., 2017), in organoids the primary cilia were not completely disassembled before mitosis. To further confirm the apical localization of the cilia, 3D reconstructions of the apical end foot of NPCs were generated, which revealed the presence of a single cilia pointing towards the lumen in organoids (Figure 3 E’’). These observations demonstrate that P and A organoids exhibit the characteristic presence and localization of primary cilia, similar to the NPCs in the developing NT.

During NT formation, the apical membrane of NPCs undergoes reorganization into distinct micro-domains. In these micro-domains, specific proteins involved in junctional complexes (such as N-cadherin, α-catenin, and β-catenin) are located in the subapical domain, zonular proteins (including ZO1, afadin, and actin) occupy an intermediate position, and fate-determining factors (such as PAR3, aPKC, and prominin-1) are confined to the most external domain(Aaku-Saraste et al., 1996; Afonso and Henrique, 2006; Chenn et al., 1998; Marthiens and ffrench-Constant, 2009). Transversal and *en face* imaging of organoids revealed the localization of N-Cadherin and β-Catenin at the apical belt of NPCs, resembling the distribution observed *in vivo* (Figure 3G and Supplementary Figure 4E,F). Furthermore, co-staining of aPKC and ZO1 demonstrated that NPCs within both A and P organoids organize membrane microdomains lining the lumen, closely resembling the organization seen in NPCs of the developing NT. In this arrangement, the aPKC microdomain faces the organoid lumen and is positioned external to the ZO1 membrane domain (Figure 3 G,H). These analyses collectively indicate that both A and P human organoids are composed of NPCs exhibiting a cell and tissue architecture highly similar to that of the early embryonic NT.

To directly investigate the dynamics of neural progenitor cells (NPCs) within the human organoids we employed the piggyBac™ transposon system (pCAG-PBase) to electroporate hESCs with a GFP vector (PBCAG-eGFP). The GFP-expressing hESCs were cultured under conditions that guide the formation of mosaic posterior neural organoids, which were then imaged using an Andor Dragonfly 505 confocal microscope. This imaging system allowed organoids to grow at a similar rate as in normal culture conditions and enabled the segmentation and tracking of fluorescently labelled electroporated cells over time (Figure 4 A, B). Time-lapse imaging of human organoids revealed the stereotypic movement pattern of NPCs synchronized with the cell cycle, known as INM ^61^. GFP+ cells in contact with the organoid ventricle displayed a high circularity associated with the rounded nucleus during mitosis (M) (Figure 4 C and Supp. Movies S1, S2). Tracking the GFP+ cells allowed us to estimate the total cell cycle length within a span of 40 hours by measuring the time taken between cell shape changes, as well as the time taken for cells to transit from the organoid lumen to the basal end and back to the lumen (Figure 4D, and Supp. Movies S1, S2). Throughout the cell cycle, the nucleus of NPCs exhibited specific movements: during the G1 phase, the nucleus travelled basally, and the cell extended two feet contacting the apical and basal tissue limits; during the S phase, the nucleus contacted the basal region (periphery of the organoid); and during the G2 phase, the nucleus travelled apically toward the organoid lumen, while the cell extended two feet contacting the apical and basal regions (Figure 4 E and Supp. Movies S1, S2). Additionally, cell tracking enabled the estimation of the duration of different cell cycle phases: G1 phase (median ±IQR time = 2.25 ±5.12 hours), S phase (median ±IQR time = 26.75 ±7 hours), G2 phase (median ±IQR time = 9.5 ±3.63 hours), and M phase (median ±IQR time = 1 ±0.38 hours) (Figure 4D and Supp. Movies S1, S2).

**Figure 4:**
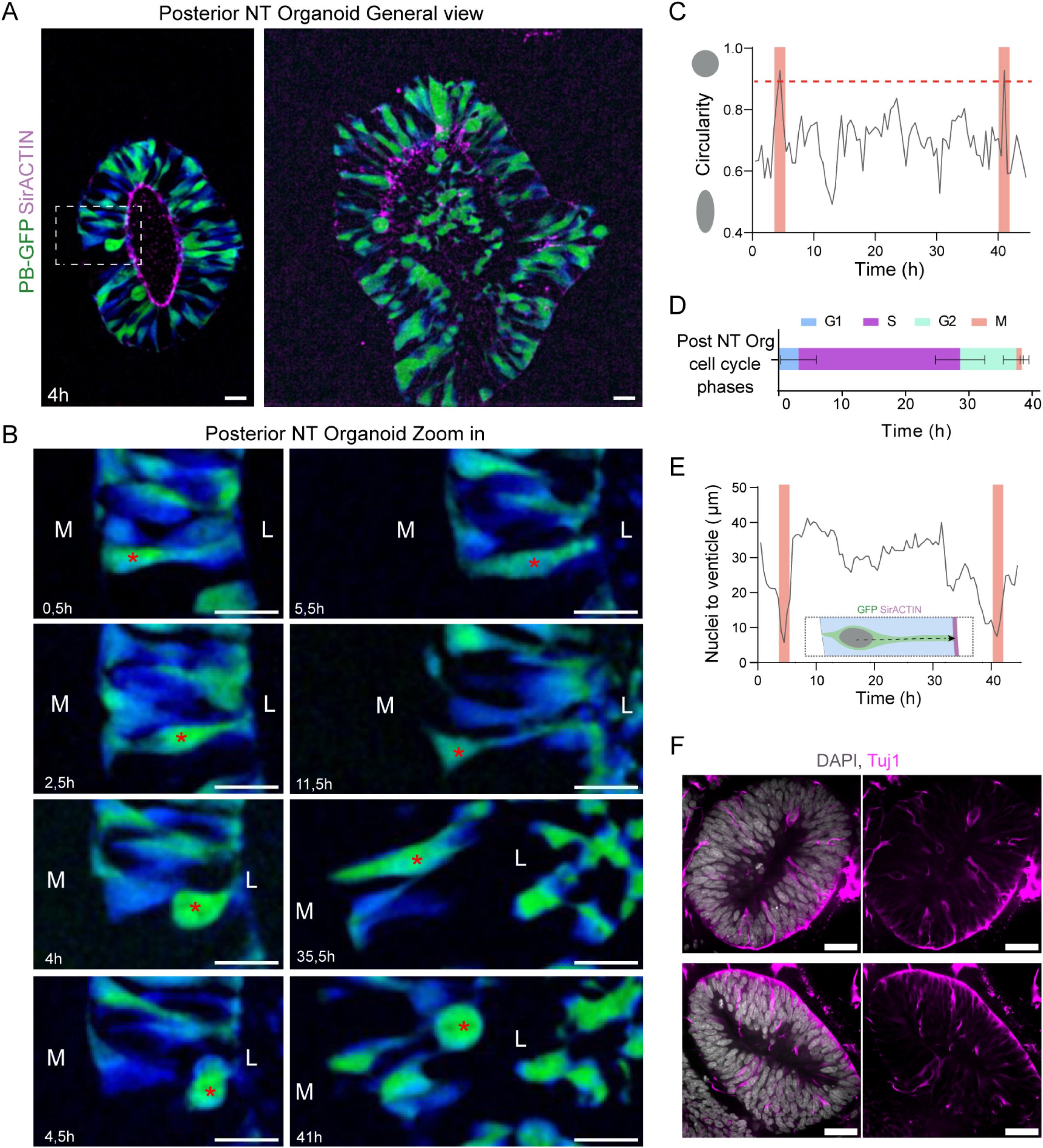
The cell and tissue dynamics of neural progenitor cells within the human organoids mimic the early embryonic NT. **(A)** Selected images of GFP-expressing posterior organoids, at the beginning and the end of the time-lapse video microscopy, where GFP+ cells are tracked undergoing INM. *In vivo*, organoids are stained for ACTIN (magenta) and GFP (green to blue). Scale bars = 10 µm. **(N)** Plots the cell cycle length estimation from the time-lapses (n=5). **(B)** Zoom in of the sequential time point following a NPC (marked with the red asterisk) undergoing INM. NT organoids are labelled for GFP (green to blue). L= Lumen; M= Matrigel. Scale bars = 10 µm. **(C)** Plot shows the changes in circularity that the NPC undergoes in the time-lapse showed in **B**. Orange bars highlight the mitosis time points. **(D)** Plots the cell cycle length estimation from the time-lapses (n=5). **(E)** Plots the position of the cell nuclei within the organoid, as the distance to the lumen surface in the time-lapse showed in B. Orange bars highlight the mitosis time points. **(F)** Selected images of posterior NT organoids, 10 days in culture showing laterally positioned Tuj1 (magenta) expressing differentiated neurons, DAPI (white). Scale bar= 10 µm.

Neurogenesis in the embryonic NT requires the detachment of newly born neurons from the neuroepithelium, referred to as delamination. This process plays a key role in ensuring the correct positioning of neurons, which is essential for shaping neuronal architecture and establishing functional circuitry(Saade and Martí, 2025). Results show that, cells expressing pan-neuronal markers such as Tuj1, detach from the organoid lumen, delaminate laterally to occupy the periphery of the organoid, mimicking the behavior of primary neurogenesis in the embryonic spinal cord(Saade and Martí, 2025)(Figure 4 F).

Overall, these data demonstrated the robustness of the *in vitro* generated neural organoids, not only in terms of transcriptional identity but also in terms of cellular architecture and dynamics. Consequently, these unique biomodels can be utilized to investigate the biological processes involved in posterior spinal cord development, which is poorly understood but crucial as abnormalities during this process can lead to neural tube defects, among the most common birth defects observed in humans.

### Modelling secondary neurulation in human spinal cord organoids

Our current understanding of how the posterior spinal cord develops in human embryos is fragmentary(O’Rahilly and Müller, 1987; Saitsu et al., 2004; Saitsu and Shiota, 2008; Santos et al., 2023), and it is largely based on studies in vertebrate animal models. In the chick embryo, we have shown that the confinement of NMPs into the prospective medullary cord (which is the immediate precursor of the secondary NT, or SNT) is concomitant with the deposition of an extracellular matrix (ECM) containing classical cell-surface-associated glycoproteins(Gonzalez-Gobartt et al., 2021) As NMPs transform into fully epithelialized NPCs, multiple small lumens emerge *de novo* at the interface between the peripheral epithelial and central mesenchymal cell populations. These small lumen foci appear at a distance equivalent to one cell’s length from the developing basement membrane(Gonzalez-Gobartt et al., 2021)

In line with this *in vivo* process, our study demonstrated that hESCs possess an intrinsic capacity to self-polarize and self-organize around a single central lumen. However, it should be noted that our culture conditions involve the clonal growth of small clusters of hESCs, where all the cells are in contact with a matrigel basement membrane-like environment. As a result, the forming organoids lack central cells with mesenchymal polarity (Figure 1 B, C). To better mimic the *in vivo* secondary neurulation cell rearrangements, we hypothesized that starting with a larger clump of hESCs would allow the presence of central cells isolated from the matrigel. Hence, we maintained GFP-expressing hESCs in agitation for 48 hours to form cell aggregates with mesenchymal features (Figure 5 A, B). After transferring these aggregates into 3D matrigel and culturing them in neural induction medium supplemented with WNT/bFGF, BMP inhibition, and RA, neural organoids were formed. In these organoids, peripheral cells elongated and acquired epithelial polarity while central cells retained mesenchymal morphology, resembling the embryonic medullary cord (Figure 5 B). Importantly, under these culture conditions, de novo emergence of small lumen foci occurred at a distance equivalent to one cell’s length from the matrigel. Over time, these small foci resolved into a single central lumen (Figure 5 B), closely resembling the morphogenetic events observed during posterior spinal cord development in both humans and chick embryos through secondary neurulation(Gonzalez-Gobartt et al., 2021; O’Rahilly and Müller, 1987; Saitsu et al., 2004; Saitsu and Shiota, 2008; Santos et al., 2023).

**Figure 5:**
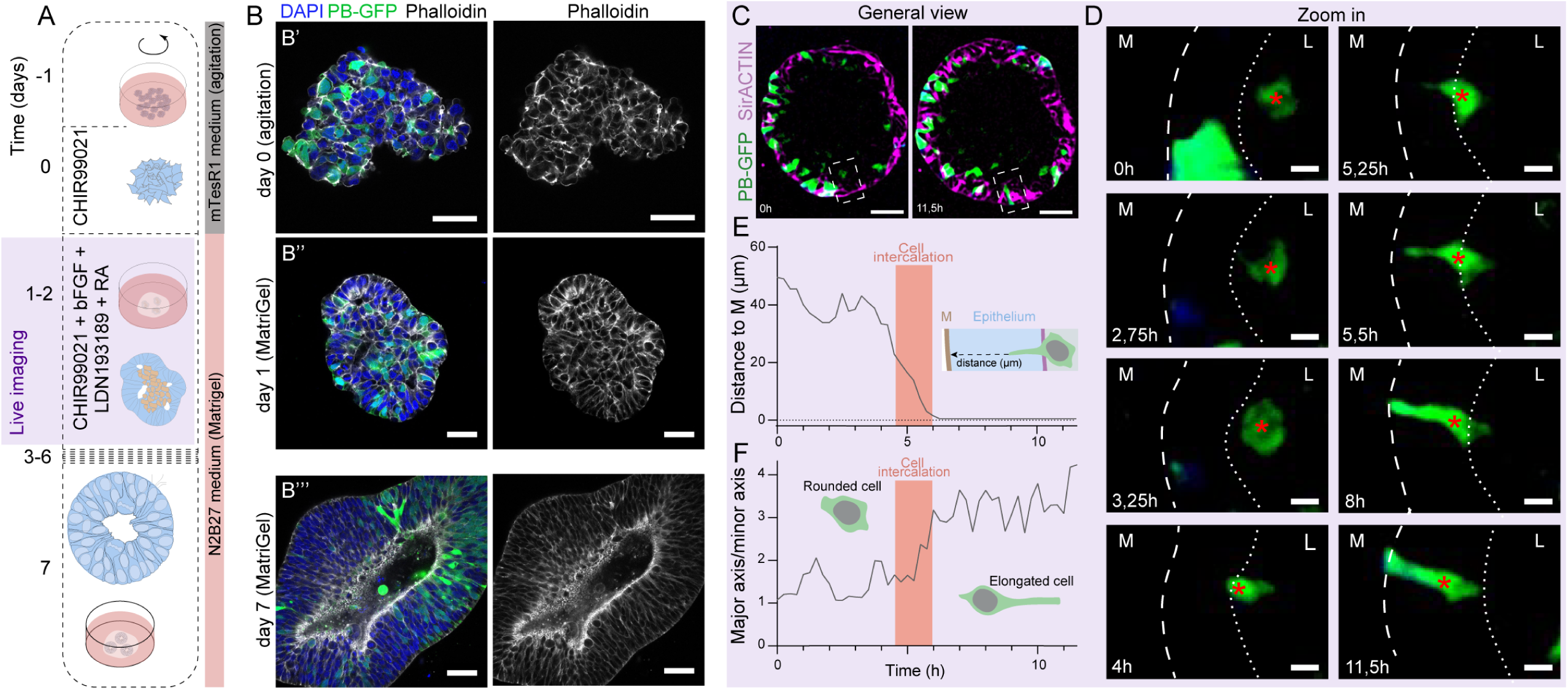
Modelling *de novo* lumen formation in posterior organoids and tracking cell intercalation for lumen resolution. **(A)** Scheme representing the pre-aggregation step protocol for modelling secondary neurulation in posterior NT organoids. The violet square represents the live imaging time window showed in C and D. **(B)** Selected images of the general view of posterior organoids maintained 48h in agitation and then transferred to a Matrigel droplet. **B’’** shows a selected image of posterior organoids undergoing multiple lumen formation that spontaneously will later resolve into a single lumen **(B’’’)**. Organoids are stained for ACTIN (grey), GFP (green) and DAPI (blue). Scale bars = 50 µm. **(C)** Selected images of the general view of posterior organoids at the beginning and the end of the time-lapse. Organoids are stained for ACTIN (magenta) and GFP (green to blue). Scale bars = 30 µm. **(D)** Zoom in of the sequential time point following a NPCs (marked with the red asterisk) undergoing cell intercalation in the posterior organoids showed in C. Taking the ACTIN staining as reference, a long discontinuous line was drawn to mark the BM while the short discontinuous line represents the apical limit of the epithelia. L= Lumen; M= Matrigel. Scale bars = 10 µm. **(E)** Plot shows the minimum distance of the intercalating cell to the BM showed in D as time goes on. Pink areas highlight the intercalation time points. **(F)** Plots the changes in cell shape assessed by the ratio between major and minor axis of the intercalating cell showed in D as time goes on. Pink areas highlight the intercalation time points.

Notably, this capacity to organize a solid medullary cord and to open multilumens is restricted to the Posterior neural organoids. Results show that hESC that were maintained in agitation for 48 hours to form cell aggregates with mesenchymal features, when guided to form Anterior neural organoids by a 48-hour pulse of TGFβ/BMP inhibition, rarely organize multiple lumens (Supplementary Figure 5A-F). The differentially expressed genes (DEGs) identified when comparing the transcriptomes of A and P organoids (Figure 2C), might be dictating these different tissue behaviours further indicating that, this is a morphogenetic property associated to the transcriptional identity of posterior spinal cord organoids, that indeed mimics the morphogenesis of the posterior secondary NT of the elongating embryo.

In the chick embryo posterior spinal cord, the resolution into a single lumen involves the intercalation of centrally located cells among the epithelialized NPCs, a process independent of apoptosis(Gonzalez-Gobartt et al., 2021). Our analysis of the human posterior NT organoids aimed to test the conservation of these mechanisms. We tracked the dynamics of GFP-expressing cells within the posterior NT organoids using time-lapse microscopy (Figure 5 C). Observations of GFP+ cells revealed the presence of round mesenchymal cells located centrally within the organoids (indicated by a red asterisk and depicted in the initial time-points in Figure 5 D). Importantly, we consistently observed these central cells intercalating into the lateral walls of the neuroepithelium, resulting in a lateral movement away from the centre of the organoid (Figure 5 D; Supp. Movies S3-6). Notably, the intercalating cells exhibited a round shape and displayed high protrusiveness. The intercalation of these cells consistently led to cell elongation (Figure 5 E, F; Supp. Movies S3-6). These findings suggest that the intercalation of centrally located cells into the lateral walls of the neuroepithelium, resulting in cell elongation, is a conserved mechanism for the resolution into a single continuous lumen in the posterior human NT organoids, similar to what is observed in the chick embryo secondary neural tube formation. Importantly, this process does not involve apoptosis, as demonstrated by the absence of cleaved caspase-3 staining in the peripheral NPCs or the inner cells surrounded by forming lumens in the organoids (Supplementary Figure 5 G-I).

### Lumen resolution in posterior neural organoids involves YAP mediated cell intercalation

In our analysis of posterior NT organoids, we made an intriguing observation regarding cell density during the process of cell intercalation. We found that cell density shifted from lower levels in the central regions to higher density in the peripheral NPCs, with cells occupying larger areas in the center and becoming more compact toward the periphery (Supplementary Figure 6A,B). The Hippo effectors YAP and TAZ function as mechanosensitive on–off switches that respond to changes in extracellular matrix (ECM) composition and mechanics. Accordingly, increased cell density is known to suppress Hippo pathway activity, thereby promoting the nuclear translocation of its downstream effector, YAP(Elosegui-Artola et al., 2017; Nardone et al., 2017; Varelas et al., 2010, 2008).

To investigate the role of YAP in lumen resolution, we examined endogenous YAP protein levels in Posterior human neural organoids during the clearance of central mesenchymal cells (Figure 6A). Immunostaining for YAP revealed that active YAP, assessed by the nuclear-to-cytoplasmic ratio, was higher in the central intercalating cells than in the peripheral NPCs (median ± IQR nuclear/cytoplasmic YAP ratio in central cells = 0.43 ± 0.20 vs. peripheral cells = 0.30 ± 0.11; Figure 6B). This pattern of YAP activity correlates with the observed shift in cell density—from lower density in the central region to higher density in the peripheral NPCs (median ± IQR cell area in central cells = 178 ± 66.3 µm² vs. peripheral cells = 115 ± 37.8 µm²; Figure 6C). These findings suggest a conserved role for YAP activity in regulating cell intercalation and lumen resolution in human organoids, consistent with previous observations in the chick embryo, where YAP activity governs lumen resolution during secondary neurulation(Gonzalez-Gobartt et al., 2021)

**Figure 6:**
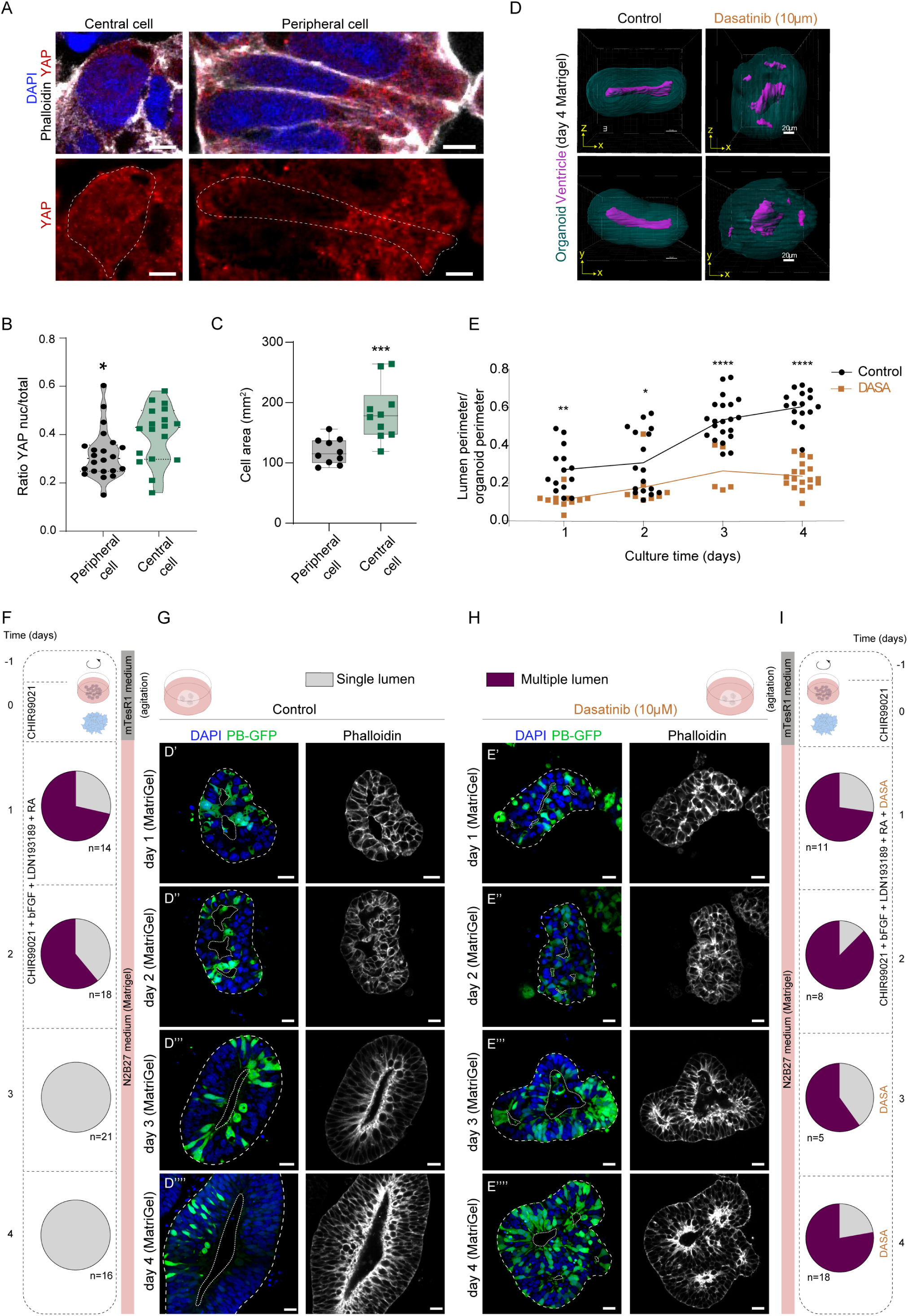
Lumen resolution in posterior neural organoids requires YAP activity. **(A)** Selected images of peripheral and central cells from Posterior organoids maintained 48h in agitation and then grown in a Matrigel droplet for 24h. Organoids are stained for ACTIN (grey), YAP (red) and DAPI (blue). Dot line represents the limits of the cells based on the ACTIN staining. Scale bars = 5 µm **(B)** Plots the ratio of the integrated density of the nuclear YAP over the integrated density of YAP in the whole cell (horizontal bold lines show the median; n=23 (peripheral cells) n=18 (central cells), from 7 organoids; *p<0.05 Mann-Whitney test). **(C)** Plots the mean area occupied by peripheral and central cells (horizontal bold lines show the median; n=10, 10 images from 5 organoids; ***p<0.001 Mann-Whitney test). **(D)** 3D reconstructions of organoid lumen (ventricle) in control and Dasatinib (DASA) treated cultures. **(E)** Plots the ratio of lumen perimeter mean over the organoid perimeter at day 1, day 2, day 3 and day 4 of culture in control (black) and DASA (orange) (horizontal bold lines show the median; * p<0.05, ** p<0.005, **** p<0.0001, two-way ANOVA). **(F)** Schematic representation of the culture conditions Pie charts represent the proportion of organoids with single (grey) and multiple (purple) lumen. **(G,H)** Selected images of the general view of posterior organoids maintained 48h in agitation and then transferred to a Matrigel droplet in control conditions or with a YAP inhibitor (DASA). Organoids are stained for ACTIN (grey), GFP (green) and DAPI (blue). Scale bars = 20 µm. White dot line delimitates the lumen perimeter while, the white discontinuous line delimitates the organoid perimeter. **(I)** Schematic representation of the culture conditions. Pie charts represent the proportion of organoids with single (grey) and multiple (purple) lumen.

To further investigate the requirement of YAP activity in cell intercalation and lumen resolution, we conducted experiments using dasatinib, a C-SRC/YAP1 inhibitor(Hsu et al., 2018). 3D image reconstruction of NT organoids showed that after 4 days in matrigel, the control organoids resolve a single central lumen, while dasatinib -treated organoids present a multilumen phenotype (Figure 6 D, E; Supp. Movies S7 and 8). Following the secondary neurulation morphogenic progression, we observed that control organoids after 3 days of culture in matrigel had already coalesced the lumen foci into a single central lumen (Figure 6 F, G; day 2 7/18, day3 21/21, day4 16/16). Conversely, the organoids treated with dasatinib were unable to resolve a single lumen and instead exhibited a multi-lumen phenotype even after 4 days of culture, resembling neural tube defects (Figure 6 H, I; day 2 1/8, day3 2/5, day4 4/18). This indicates the crucial role of YAP activity in the proper resolution of lumens and suggests that dysregulation of YAP signalling could contribute to neural tube defects such as spinal bifida.

These findings demonstrate that the Posterior human NT organoids not only exhibit a transcriptome profile resembling the developing spinal cord of the early human embryo, but also replicate cellular dynamics observed during the morphogenesis of the human embryo’s secondary neural tube. This biomodel provides a unique platform to directly study the underlying mechanisms of neural tube defects such as spinal bifida.

## DISCUSSION

In this study, we successfully replicated the early morphogenesis of the posterior human spinal cord using *in vitro* techniques. We directed human pluripotent stem cells, including both embryonic stem cells and induced pluripotent stem cells, to differentiate into neural cells, specifically those with a spinal cord identity. By creating a culture environment that mimicked the conditions of the elongating embryo, we generated 3D organoids that exhibited a transcriptional profile indicative of spinal cord tissue. These spinal cord organoids were composed of neuroepithelial cells that displayed a subcellular architecture and dynamics similar to that of the early embryonic NT, as well as reproducing the transcriptome identity of the early embryo spinal cord (5 weeks of development). Numerous organoids have been developed that mimic several traits of NT development(Gouti et al., 2017, 2014; Gribaudo et al., 2024; Karzbrun et al., 2021; Lee et al., 2022; Libby et al., 2021; Rito et al., 2025; Xue et al., 2024); nonetheless, compared to previous, the protocol described in this study is shorter. Moreover, the cellular dynamics observed in the organoids also resembled the morphogenetic events involved in secondary neurulation, a process poorly understood, hence these models might bridge the knowledge gap regarding the formation of the human posterior spinal cord.

This understanding is crucial for elucidating the biological bases of neural tube defects (NTDs) which are frequent congenital disorders, thereby providing significant medical insights into primary prevention and reduce suffering caused by loss of movement in the legs, and bladder and bowel dysfunction, caused by NTDs such as spina bifida. Even though NTDs are common congenital malformations, their causes remain largely unknown. Environmental risks, such as folic acid deficiency and maternal diabetes(Blom et al., 2006; McNeely and Howes, 2004) yet do not account for the estimated heritability of 60–70%. A recent large-scale study identified 187 genes with damaging de novo mutations in individuals with spina bifida (Ha et al., 2025). These genes cluster in developmental pathways, including Wnt signalling, planar cell polarity (PCP), and cytoskeleton regulation(Ha et al., 2025), but the cell- and tissue-level consequences remain unclear. Two further studies highlighted structural variants as genetic risk factors: for 22q11.2 deletions(Vong et al., 2024) and rare copy number variations in genes associated with metabolic pathways and the WNT/ PCP pathway(Wolujewicz et al., 2021). These genetic data, together with our experimental analysis in the chick embryo secondary neurulation, support the hypothesis that NTDs might arise from defects in the NMPC dynamics, such as a) the mesenchymal-to-epithelial transition of NMPCs required for full epithelialization of progenitor cells in building the caudal NT, and b) the tissue rearrangements required for the de novo lumen formation and single lumen resolution, to build a hollow NT. These are morphogenetic processes that are still to be revealed in human development, and the organoids engineered in this study holds promise for advancing our understanding of the biological bases and the tissue dynamics associated with secondary neurulation.

Our previous study conducted in chick embryos revealed the involvement of YAP/TAZ(Gonzalez-Gobartt et al., 2021), which act as key sensors of cell density(Piccolo et al., 2014), in the intercalation of central cells during the resolution of the lumen in the secondary NT Moreover, the organization of the actin cytoskeleton and the activation of YAP are required and play differential roles in modulating different phases of neural organoids self-organization(Tang et al., 2022). Consistent with this model, human spinal cord organoids generated in this study exhibited higher levels of nuclear (active) YAP in the central cells compared to the peripheral neuroepithelial cells. Furthermore, when the Hippo pathway, which regulates YAP/TAZ activity, was blocked using the pharmacological inhibitor Dasatinib, the process of lumen resolution was impaired, resulting in a multilumen phenotype resembling NTDs. These findings support the idea that the YAP pathway mediates lumen resolution in response to cell density during secondary neurulation. A mechano-transduction role for YAP in NT formation has been proposed in mouse embryos, since the surface ectoderm exhibits spatially heterogenous tension that correlates with YAP localisation during spinal NT closure(Marshall et al., 2023). In addition, YAP regulates expression of key patterning factors (FoxA2, Shh), and can produce NTDs(Cheng et al., 2023).

While further research is necessary to fully understand the mechanisms underlying secondary neurulation and its association with NTDs, the results of this study lay the groundwork for generating patient-derived organoids. This approach holds promise for advancing our understanding of diseases associated with secondary neurulation and may contribute to the development of improved therapeutic strategies including the use of YAP/TAZ activators to alleviate some NTDs.

## MATERIALS AND METHODS

### EXPERIMENTAL MODEL AND SUBJECT DETAILS

#### hESCs Culture

Female RUES2 (Rockefeller University Embryonic Stem Cell Lines; NIH Registration number 0013, WiCell, RRID: CVCL_B810) human ES cells isolated from the inner cell mass of human blastocysts were cultured on Matrigel coated tissue culture dish with mTeSR1 media in a humidified 37°C incubator with 5% CO2. Cells were passaged every 3-5 days as follows:

- Cells were rinsed 2 x in PBS (Fisher Scientific #10010015).
- Cells were incubated in PBS-EDTA 0.5mM (Sigma #20-158) for 4 min at 37°C.
- Detached multicellular clumps were collected in mTeSR1 (StemCell #85850) and diluted between 1/4 and 1/10.
- Clumps were transferred to a six-well tissue culture plate (Corning #353046) (6WP) precoated with 1% lactate dehydrogenase–elevating virus (LDEV)–free hESC-qualified BM matrix Matrigel (Corning #354277) for 30 min to 2h.

All cell lines used in this study had a passage number of <P100. They were tested negative for mycoplasma contamination.

All experiments involving hESCs were approved by CSIC Ethical Committee protocol code 089/2022

#### hiPSC Culture

Male CRTD1 (CRTDi004-A) (Centre for Regenerative Therapies Dresden (CRTD), hPSCreg, RRID: CVCL_YR23) and KOLF2C1 (WTSIi018-B-1) (Welcome Sanger Institute (WTSI), hPSCreg, RRID: CVCL_9S58) human iPS cells were cultured on Matrigel coated tissue culture dish with mTeSR1 media in a humidified 37°C incubator with 5% CO2. We did not authenticate CRTD1 and KOLF2C1 cell lines. Cells were passaged every 3-5 days as follows:

- Cells were rinsed 2 x in PBS (Fisher Scientific #10010015).
- Cells were incubated in ReLeSR (StemCell #05872) for 1 min at RT.
- Detached multicellular clumps were collected in mTeSR1 and diluted between 1/4 and 1/10.
- Clumps were transferred to a six-well tissue culture plate (Corning #353046) (6WP) precoated with 1% lactate dehydrogenase–elevating virus (LDEV)–free hESC-qualified BM matrix Matrigel (Corning #354277) for 30 min to 2h.

All cell lines used in this study had a passage number of <P100. They were tested negative for mycoplasma contamination.

All experiments involving hiPSCs were approved by CSIC Ethical Committee protocol code 089/2022

### METHODS DETAILS

#### hESC electroporation

The piggyBac™ transposon system was used to integrate a GFP sequence into the genome of the hESC. piggyBac transposase vector (pCAG-PBase) was co-nucleofected with a piggyback transposon GFP vector (PBCAG-eGFP) (Lacoste et al., 2009) to generate hESC mosaic cultures that ultimately are used to generate mosaic organoids with GFP+ and GFP-cells that allow a better characterization of cell rearrangements along the secondary neurulation process and organoid *in vivo* imaging. Nucleofection of undifferentiated hESC were carried out as follows:

- Cells were rinsed 2 x in PBS.
- Cells were incubated in StemPro™ Accutase™ (Thermo Fisher #A1110501) for 3 min at 37°C.
- Cell suspension was recovered in 3mL of mTesR1 with Y27632 2HCl 10μM (Selleckchem #S1049).
- Cell suspension was spin down and cell density was quantified.
- 8 x 105 cells were resuspended in 100μL of Human Stem Cell Nucleofector® Solution (Lonza # VPH-5002) with 4μg of PBase and 4μg of PB-GFP DNA vectors.
- Solution is transferred to an electroporation cuvette (Lonza # VPH-5002) to subsequently been inserted into the Nucleofector® 2b Device (Lonza #AAB-1001) to apply program A-012.
- Cell were recovered in mTesR1 with Y27632 2HCl 10μM to subsequently being seeded in 2D for maintenance or directly to generate matrigel droplets and undergo organoid guidance.

#### Nerual organoids differentiation guidance

Most of the organoid experiments were carried out with the RUES2 cell line. However, there were no significant differences when organoid differentiation was done with the other cell lines. Guidance went as follows:

- Cells were rinsed 2 x in PBS.
- Cells were incubated in StemPro™ Accutase™ for 3 min at 37°C.
- Cell suspension was recovered in 3mL of mTesR1 with Y27632 2HCl 10μM.
- Cell suspension was spin down and cell density adjusted to 10 x 10^6^ cells/mL.
- 10^5^ cells were incorporated to 150μL of Matrigel on ice.
- A droplet of 10uL of the Matrigel cell suspension was pipetted per well in a twenty-four-well tissue culture plate (Corning #353046) (24WP).
- Droplet plates were incubated for 13 min at 37°C to subsequently add 0,5mL of mTeSR1 with Y27632 2HCl 10μM.

The following days mTeSR1 is substituted by N2B27-based neural induction medium comprised of: Advance DMEM/F12 (Gibco #31331-028):Neurobasal medium (1:1; Gibco #21103-049), 0,5× N2 (Gibco #17502-001), 0,5× B27 (Gibco #17504-001), 1×nonessential amino acids (Gibco #11140-035), 1x Sodium Piruvate (Gibco #11360-088), 0,5x GlutaMax (Gibco #35050-038), and 0,1 mM β-mercaptoethanol (Gibco #31350-010). Media was supplemented with the appropriate drugs as indicated in each experiment. For references and concentrations of the drugs see KEY RESOURCES TABLE. Media was changed every day and plates were kept at 37°C and 5% of CO_2_.

#### Pre-aggregation organoid protocol

To study the epithelialization and lumen resolution processes, solid aggregates were generated before the Matrigel induction process. This was achieved with a two-day spheroid preassembling protocol that was carried out as follows:

- Cells were rinsed 2 x in PBS.
- Cells were incubated in PBS-EDTA 0.5mM for 4 min at 37°C.
- Detached multicellular clumps were collected in mTeSR1.
- Approximately 10^6^ cells forming clumps were seeded in a 6WP with mTesR1 and kept shaking at 150rpm at 37°C and 5%CO2.
- 3μM CHIR99021 (Milipore # 361571) was added to the media the following day.
- One day after, spheroids from each well were collected, spin down and re-suspended between 50-80μL of Matrigel to generate droplets in 24WPs and complete the organoid guidance as it is detailed in the previous section.

#### In droplet organoids immunohistochemistry

Immunostaining of organoids droplets was carried out as follows:

- Media was removed and organoids droplets were rinsed 2 with PBS.
- Organoids droplets were fixed in 4% PFA (Sigma #30525-89-4) in 1xPBS for 1 hour at RT.
- Organoids droplets were rinsed 3 x with PBS.
- Organoids droplets were permeabilized 1-3h in PBS SDS 0.1% (Merck #151-21-3).
- Organoids droplets were rinsed 3 x with PBS.
- Organoids droplets were incubated in blocking solution (10% BSA (Merck #9048-46-8) in PBT) overnight at 4°C.
- With a metal spatula organoids droplets were detached from the bottom of the 24WP and transferred to Eppendorf tubes (Cultek #F7TUBE1EP5-S).
- Organoids droplets were incubated in antibody solution (1% BSA in PBT) with primary antibody overnight at 4°C in rotation.
- Following incubation, organoids droplets were washed 3 x 5 min with PBS.
- Organoids droplets were then incubated in secondary antibodies in antibody solution overnight at 4°C in rotation.
- Finally, organoids droplets were initially washed 3 x 10 min in PBS washes. Each droplet was then transferred to a porta with a μwell sticker (iSpacer 0.2mm SUNJin Lab #IS016) where liquid is removed as much as possible and RapiClear 1.52 (SUNJin Lab #RC147001) is added on top. Finally, a glass coverslip is placed on top.

#### Free-floating organoids immunohistochemistry

Alternatively, organoids can be removed from the matrigel before the fixation and the immunohistochemistry. The free-floating organoids immunostaining was carried out as follows:

- Media was removed and organoids droplets were rinsed with PBS.
- Organoid droplets were incubated with Collagenase IV (3mg/mL) (Thermo Fisher Scientific #17104019) and Dispase (1mg/mL) (Thermo Fisher Scientific #17105041) in HBSS (Gibco # 14025-050) at 37°C and 5% CO2 for 30 mins with gentle agitation.
- Matrigel dissolution was facilitated with up and down pipetting. Finally, organoids were collected in N2B27 and spin down to fix in PFA 4% (Sigma #30525-89-4).
- Immunostaining procedure goes as has been described above for the droplets.
- Free-floating organoids were spin down and resuspended directly in RapiClear

1.52 (SUNJin Lab #RC147001). Then they were transfer to a porta with a μwell sticker (iSpacer 0.2mm) and a glass coverslip is placed on top.

#### RNA extraction from organoids

Anterior and posterior huNTOrg were extracted from the matrigel droplet as described in the previous section, pulled down together, and total RNA was extracted using the TRIzol RNA Isolation Reagent (Thermo Fisher Scientific #15596026) according to the manufacturer’s protocol.

#### Data preprocessing and RNA-sequencing analysis of Anterio and Posterior organoids samples

Bulk RNA-sequencing of Anterior and Posterior organoid samples was performed at BGI-Shenzhen using the DNBSeq-G400 platform. Quality control of the raw paired-end FASTQ files was conducted using FASTQC v0.11.9 **(1)** software. Sequence reads were trimmed for adaptor/low-quality sequence using SOAPnuke v2.1.7 (parameters: -l 15 -q 0.2 -n 0.05) **(2)**. The processed FASTQ files were mapped to GRCh38.p13 using HISAT2 v2.1.0 (parameters: --sensitive –no-discordant --no-mixed -I 1 -X 1000 -p 8 --rna-strandness RF) **(3)**. FeatureCounts v2.0.3 **(4)** was used to quantify the number of reads assigned to each gene annotated in the Ensembl GRCh38 human annotation file from release 108 (parameters: -p --countReadPairs -t exon -g gene_id -O -T 8).

#### Statistical analysis

A tab-delimited text file including raw count values was used as input matrix for the statistical analysis. DESeq2 v.1.41.1 **(5)** was used to normalize the raw data applying the median of ratios method and to identify differentially expressed genes between Anterior and Posterior organoids using a 5% false discovery rate. The correlation heatmap was drawn calculating the Pearson correlation coefficients of all gene expression between each two samples. The correlation coefficients were associated to a colour scale where darker colours represented higher correlations and lighter colour represented lower correlations. The volcano plot representation was drawn using the genes fold change data converted to log2 and the significance values converted to log10.

#### Comparison of organoids with human embryonic samples

The paired-end RNA-sequencing data from forebrain, midbrain, hindbrain, and spinal cord samples of human embryos at Carnegie Stage 14 (E-MTAB-4840) was obtained from the HDBR database ^6^. We selected the gene expression profiles of 56 key anterior-posterior axis markers and performed a Principal Component Analysis (PCA) on the normalized (log2(TPM+1) and centered neural organoids and human samples data. For this comparative analysis, the data was centered on the average expression of Anterior organoid samples. The PCA displays the transcriptional deviations of the other organoid and human samples from the Anterior control group.

For the heatmap, the TPM datasets were log2-transformed and normalized to a Z-score across each gene. The pheatmap package (v.1.0.12) (https://doi.org/10.32614/CRAN.package.pheatmap) was used to create the heatmap, hierarchically clustering the samples using Euclidean distance.

To evaluate the neural identity of the nerual organoids, we examined the gene expression profiles of key neural, mesodermal, and endodermal markers. The TPM data was transformed by log10. The transformed data was used to create a bubble plot (ggplot2) where the size of each point is proportional to expression level in log10 of each gene per sample.

List of runs for each sample

| Run | Developmental week | Developmental Stage | Organism part |
| --- | --- | --- | --- |
| ERR1473333 | 5 | Carnegie Stage 14 | Forebrain |
| ERR1473332 | 5 | Carnegie Stage 14 | Forebrain |
| ERR1473373 | 5 | Carnegie Stage 14 | Hindbrain |
| ERR1473374 | 5 | Carnegie Stage 14 | Hindbrain |
| ERR1473437 | 5 | Carnegie Stage 14 | Midbrain |
| ERR1473438 | 5 | Carnegie Stage 14 | Midbrain |
| ERR1473478 | 5 | Carnegie Stage 14 | Spinal cord |

#### Image Acquisition of fixed samples

##### Confocal microscopy

Samples were imaged on an inverted Zeiss LSM-780 confocal microscope, equipped with an Argon multiline gas laser at 488 nm, a DPSS laser at 561 nm and a HeNe laser at 633nm. The objectives used were a ZEISS 40X (oil, NA 1.3, 0.2mm working distance) and a ZEISS 63X (oil, NA 1.4, 0.18mm working distance).

##### Fluorescence microscopy

Samples were imaged on an inverted Leica DMI8 microscope equipped with Spectra-X Light Engine: 6 solid-state LEDs. The objective used was a LEICA (oil, NA 1.4/0.7, 0.09mm working distance).

Instant computational clearing algorithm were used in image post-processing to remove out-of-focus light in real-time to enhance optical sectioning.

#### Image Acquisition of *in vivo* samples

##### Mounting

One organoid droplet was seeded per bottom-plastic dish (ibidi μ-Dish 35mm, high; #81156) as described in organoid differentiation guidance or Spheroid preassembling sections. Dishes were kept at 37°C and 5%CO_2_.

##### Staining

Organoids used for *in* vivo time-lapse imaging are a GFP+ cells mosaic, generated as explained in hESC nucleofection section.

To facilitate lumen and cell shape identication *in vivo,* 50-200nM SiR-actin (SPIROCHROME, #SC001) was added to sample media between 6 and 24 hours before imaging. It is a fluorogenic, cell permeable and highly specific probe for F-actin compatible with GFP and mCherry fluorescent proteins.

##### Spinning-disk time-lapse microscopy

Samples were imaged on an inverted Nikon Ti-E microscope stand including the Perfect Focus System (PFS) equipped with 405, 488, 561 and 637 nm laser lines. The objective used was a CFI Plan Apochromat Lambda S LWD (water, NA 1.14, 0.59mm working distance).

Images were taken every 15mins for 24 to 72h. Laser were set in low power mode at 1-2%. Image acquisition was done at high speed with a resolution of 2048×2048 and binning 3×3.

### IMAGE QUANTIFICATION AND STATISTICAL ANALYSIS

#### General image analysis

Raw confocal data was exported to ImageJ/FiJi (http://rsbweb.nih.gov/ij/) RRID: SCR_003070) to be processed and analysed. Projections of z-stacks are maximum projections unless otherwise indicated. Figures and schemes were generated using Adobe Illustrator CC2018 (RRID: SCR_014199).

3D reconstructions were generated using the 3D viewer plugin included in the ImageJ/Fiji.

#### Apoptotic index

Cleaved-Caspase3 (cCaspase3) antibody was used to detect apoptosis in fixed organoids. We counted both the number of cCapase3+ cells and the number of total DAPI cells for central cells and epithelialized cells. Percentages were then calculated and presented in GraphPad Prism 8 bar graphs (mean ± SD/s.e.m.).

#### Cell shape

βCat antibody was used to visualize cell shape in fixed organoids. Cells were delimited with the polygon selection tool and cell shape was quantified by measuring cell aspect ratio (AR), a parameter included in ImageJ Shape descriptors. AR is calculated dividing major axis diameter between minor axis diameter, with a value of 1.0 indicating a perfect circle. As the value increase over 1, it indicates an increasingly elongated shape. Results are presented in GraphPad Prism 8 violin plots.

#### Centrosome positioning

Centrosomes were visualized with PCNT antibody and DAPI was used to stain the nucleus in organoid samples. The straight line tool of ImageJ was used to draw a line from the centrosomes to the edge of the nucleus and the distance was measured. Results are presented in GraphPad Prism 8 violin plots.

#### Ciliary length

Length of the primary cilium was quantified in anti-Arl13B stained organoids. A straight or segmented line was drawn onto the Arl13B staining of each cell and length was measured with the ImageJ command. Results are represented in GraphPad Prism 8 violin plots showing all points, median and interquartile range.

#### Relative localization of proteins

ZO1 and aPKC antibodies were used to assess apical components organization. An intensity profile was generated from a line drawn along the cell’s major axis for ZO1, aPKC and DAPI. Thus, the relative apico-basal position is assessed. Results are plotted in GraphPad Prism 8 XY graphs.

Similarly, YAP antibody and Phalloidin and DAPI stainings were used to assess differences between nuclear and cytoplasmic YAP. An intensity profile was generated from a line drawn along the cell for YAP, Phalloidin and DAPI. Results are plotted in GraphPad Prism 8 XY graphs.

#### Quantification of YAP nuclear/cytoplasmic ratio

PB-GFP was electroporated in order to draw the cell shape in organoids neuroepithelial cells. DAPI staining was used to detect the nucleus. YAP was stained using a YAP antibody to assess protein levels. YAP activity was inferred from the ratio of nuclear over the cytoplasmic YAP protein levels. Nuclear regions were delimitated by hand in the ImageJ using DAPI staining as reference. Cytoplasmic region was estimated subtracting the nuclear area to the total cell area delimitated by hand in the ImageJ using the GFP staining as reference. YAP protein levels were assessed in the already defined nuclear and cytoplasmic areas through the ratio of RawIntDensity over the area extension using ImageJ/Fiji. Distance was presented in GraphPad Prism 8 violin plots.

#### 3D lumen reconstruction of neural organoids

Raw confocal data of Posterior organoids was exported to the Imaris software (Bitplane) (RRID:SCR_007370). The secondary forming lumen was reconstructed using the Contour Surface. The 3D structure was extracted by manually drawing the lumen contour, visible with aPKC immunostaining, on consecutive 2D z-slices.

#### Quantifications in time-lapse movies

##### Distance from the soma of NPCs undergoing IKNM to the ventricle

A GFP mosaic organoid was generated by PB-GFP electroporation, allowing cell tracking. A straight line was drawn in the periphery of the ventricle based on the SiR-Actin staining. Mitosis were spotted and tracked throughout the movie. The distance from the drawn midline to the soma of the analysed cell and its daughter cell was measured in each time point with the ImageJ straight-line tool. Results along time are presented in GraphPad Prism 8 linear regression graphs.

#### Circularity of NPCs undergoing IKNM

A GFP mosaic organoi was generated by PB-GFP electroporation, allowing cell tracking. Mitosis were spotted and tracked throughout the movie. Cells were delimited with the polygon selection tool of Image J and cell shape was quantified by measuring cell circularity in each time point. Circularity is calculated with the formula 4π×[Area]/[Perimeter]2, with a value of 1.0 indicating a perfect circle. As the value approaches 0.0, it indicates an increasingly elongated shape. Results along time are presented in GraphPad Prism 8 linear regression graphs.

#### Distance from centre cells to the periphery of the organoids

GFP mosaic organoids were generated by PB-GFP electroporation, allowing cell tracking. A straight line was drawn in the periphery of the organoid based on the SiR-Actin staining. Central cells were identified based on the SiR-acting staining and in its mesenchymal shape. The distance from the drawn line to the closes side of the analysed cell was measured in each time point with the ImageJ straight-line tool. Results along time are presented in GraphPad Prism 8 linear regression graphs.

#### Major/minor axis of central cells undergoing intercalation

GFP mosaic organoids were generated by PB-GFP electroporation, allowing cell tracking. Cells were delimited with the polygon selection tool of Image J and cell shape was quantified by measuring cell aspect ratio (AR), a parameter included in ImageJ Shape descriptors. AR is calculated dividing major axis diameter between minor axis diameter, with a value of 1.0 indicating a perfect circle. As the value increase over 1, it indicates an increasingly elongated shape. Results are presented in GraphPad Prism 8 linear regression graphs.

### Statistical analysis

Statistical analysis was performed using the GraphPad Prism 8 (RRID: SCR_002798). Significance was assessed by performing the Mann-Whitney test when comparing two populations or the Kruskal-Wallis when comparing more than two. In this later case, Dunn’s multiple comparisons test was also run. (*p<0.05, **p<0.01, ***p<0.001 and ****p<0.0001).

### Limitations of study

In the study, we acknowledge the presence of inter-organoid variability in the generated organoids. While we observed reduced variability compared to longer protocols for organoid generation, we still noted some variability within the droplets and cell lines used. Specifically, a small proportion of organoids did not exhibit the expected lineage restriction. This variability should be considered when utilizing the model for pharmacological screenings, and further refinement of the protocols may be necessary to improve the consistency and reliability of the organoid system.

## ACKNOWLEDGMENTS

The authors are indebted to Leica Microsystems for supporting and collaborating with the MIP Platform (IBMB). We are grateful to researchers that kindly provided DNAs and antibodies, as indicated in the reporting summary. The work in EM’s laboratory was supported by grants PID2022-139609NB-I00 and RED2022-134100-T. JBA is a recipient of a BES-2017-080050 PhD scholarship. YEM is a recipient of a PRE2022-000760 PhD scholarship. The work in JRMM laboratory was supported by grants PID2020-112566GB-I00 and RED2022-134100-T. JMR is a recipient of a Juan de la Cierva 2022-AEI and JG of a PTA 2019-AEI

## AUTHOR CONTRIBUTIONS

**JBA** developed methodology, conceived and performed most of the experiments, analysed the data, discussed the results and revised the manuscript.

**YEM, MGV and MMF** performed a significant group of experiments, analysed the data, discussed the results and revised the manuscript.

**JMR, JC and JRMM** conceived the transcriptome profiling experiments, analysed and discussed the results, wrote those sections and revised the manuscript.

**EM** supervised the study, conceived the experiments, analysed the data, discussed the results and drafted the manuscript together with JRMM.

## SUPPLEMENTARY FIGURE LEGENDS

**Supplementary Figure S1: Screening for the guidance of hESCs towards NMPs lineage and the subsequent restriction to NPCs in 3D cultures (related to Figure 1).**

**(A)** Schematic representation of organoid culture, guiding drugs are indicated at the appropriate days in culture. Selected images of Anterior organoids stained for CDX2 (Yellow) and DAPI (blue) after 3, 5 and 7 days (D). **(B, C)** Guiding drugs are indicated at the appropriate days in culture. Selected images of organoids stained for SOX2 (red), BRA (green). (**D)** Guiding drugs are indicated at the appropriate days in culture. Selected images of Posterior organoids stained for CDX2 (Yellow) and DAPI (blue) after 3, 5 and 7 days (D). **(E,F)** Guiding drugs are indicated at the appropriate days in culture. Selected images of organoids stained for SOX2 (red), BRA (green), after 3, 5 and 7 days (D). Scale bars = 30 µm.

**Supplementary Figure S2: iPSC generated neural organoids with different anterior-posterior identities (related to Figure 1).**

**(A)** Schematic representation of anterior neural organoid culture, guiding drugs are indicated at the appropriate days (D) in culture (0-2). **(B)** Quantification of organoids expressing the indicated markerş SOX2 (red), BRA (green), and CDX2 (yellow) after 5 and 7 days (D) in culture. iPSC cell lines (Kolf2C1 and CRTD1) are indicated. Guiding drugs are indicated in each column, as well as days (D) of treatments. **(C)** Schematic representation of posterior neural organoid culture, guiding drugs are indicated at the appropriate days (D) in culture (0-2). **(D)** Selected images of anterior organoids stained for SOX2 (red) and BRA (green) and DAPI (blue), after 5 and 7 days (D) in culture. (**E)** Selected images of anterior organoids stained for CDX2 (yellow) and DAPI (blue). **(F)** Selected images of posterior organoids stained for SOX2 (red) and BRA (green) and DAPI (blue), after 5 and 7 days (D) in culture. (**G)** Selected images of posterior organoids stained for CDX2 (yellow) and DAPI (blue). Scale bars = 30 µm.

**Supplementary Figure S3: RNAseq of neural organoids revealed dorsal neural progenitor identities (related to Figure 2).**

**(A)** Bubble plots the expression levels in the Anterior (blue) and Posterior (yellow) neural organoid samples, selected genes identified in the Forebrain, Midbrain, Hindbrain, and Spinal Cord of the early human embryos (week 3 of development).

**(B)** Diagram of the DV domains in the developing NT highlighting the selected genes responsible for the identity of the ventral (p3, pMN, p2-0) and the dorsal (dP6-dP1). **(C)** Heatmap of the expression of the DV domain-associated genes highlighted in **A** in spinal cord samples isolated from CS14 (week 5 of development) human embryos and Posterior neural organoids samples.

**Supplementary Figure S4: Aosterior neural organoids are composed by epithelial polarized cells (related to Figure 3)**

**(A, B)** Selected images of mesenchyme_like (A) and Anterior neural **(B)** organoids stained for the ciliary membrane marker ARL13B (green), βCAT (magenta), and DAPI (white). Scale bars = 10 µm. Higher magnifications crops are shown.

**(C, D)** Selected images of mesenchyme-like (C) and Anterior neural (D) organoids stained for the centrosome marker pericentrin (PCNT, red), the junctional complexes protein N-Cadherin (NCAD, blue), and DAPI (white). Higher magnifications are shown. Scale bars = 10 µm.

**(E, F)** Selected images of mesenchyme-like **(E)** and Anterior neural **(F)** organoids stained for the fate-determining factor aPKC (yellow) and ZO1 (cyan). Higher magnifications are shown. Scale bars = 10 µm.

**Supplementary Figure S5: Multilumen formation is associated to Posterior organoids, and single lumen resolution is independent on apoptosis (related to Figure 5).**

**(A)** Schematic representation of the culture conditions Pie charts represent the proportion of organoids with single (grey) and multiple (purple) lumen. **(B)** Selected images of the general view of Anterior organoids maintained 48h in agitation and then transferred to a Matrigel droplet, that spontaneously organize a single lumen. Organoids are stained for ACTIN (grey) and DAPI (blue). Scale bars = 50 µm. **(C)** Selected images of the general view of Posterior organoids maintained 48h in agitation and then transferred to a Matrigel droplet, that open multiple lumen foci (day 2) that spontaneously organize a single lumen (day 7). Organoids are stained for ACTIN (grey) and DAPI (blue). **(D)** Schematic representation of the culture conditions Pie charts represent the proportion of organoids with single (grey) and multiple (purple) lumen. **(E)** Plots Circularity of the cells (4π area/perimeter) in anterior (A) compared to posterior (P) organoids. As the value approaches 0,0 it indicates increasingly elongated shape. (horizontal bold lines show the median; n=100, 100 cells from 5 organoids; ****p<0.0001 Kruskal-Wallis test and Dunn’s multiple comparison test). **(F)** Plots major axis/ minor axis of the cell (aspect ratio) in anterior (A) compared to posterior (P) organoids (horizontal bold lines show the median; n=100, 100 cells from 5 organoids; ****p<0.0001 Kruskal-Wallis test and Dunn’s multiple comparison test).

**(G)** Schematic representation of the culture conditions. GFP-expressing Posterior organoids were maintained 48h in agitation and then transferred to a Matrigel droplet and fixed after 24h for staining. **(H)** Selected image of Posterior organoids stained with Phalloidin (grey), Caspase3 (red), GFP (green) and DAPI (blue). Scale bars = 10 µm. **(I)** Plot shows the percentage of Caspase positive cells in central vs peripheral cells. Horizontal bold lines show the median; n=8 organoids; son significant differences. Mann-Whitney test.

**Supplementary Figure S6. Lumen resolution in posterior neural organoids is associated with an increase in cell density and decrease in nuclear YAP. (related to Figure 6).**

**(A)** Scheme representing central and peripheral cells in relation to cell density. **(B)** Selected image of the general view of posterior organoid maintained 48h in agitation and then maintained in a Matrigel droplet for 24h. Scale bars = 20 µm. Zoom in of peripheral and central cells. Scale bars = 5 µm. Organoids are stained for ACTIN (grey), GFP (green) and DAPI (blue). **(C‘, C”)** Plots the fluorescence intensity of YAP (red line) to the distance from the organoid lumen (zero) for the central cell and the peripheral cell. Grey boxes represents the limits of the cell based on the ACTIN staining, blue box represents the nuclei localization based on DAPI staining.

## References

Aaku-Saraste E, Hellwig A, Huttner WB. 1996. Loss of occludin and functional tight junctions, but not ZO-1, during neural tube closure - Remodeling of the neuroepithelium prior to neurogenesis. Dev Biol 180:664–679. doi:10.1006/dbio.1996.0336

Afonso C, Henrique D. 2006. PAR3 acts as a molecular organizer to define the apical domain of chick neuroepithelial cells. J Cell Sci 119:4293–4304. doi:10.1242/JCS.03170

Akin ZN, Nazarali AJ. 2005. Hox genes and their candidate downstream targets in the developing central nervous system. Cell Mol Neurobiol 25:697–741. doi:10.1007/S10571-005-3971-9

Alaynick WA, Jessell TM, Pfaff SL. 2011. SnapShot: Spinal cord development. Cell 146. doi:10.1016/j.cell.2011.06.038

Basson MA, Echevarria D, Ahn CP, Sudarov A, Joyner AL, Mason IJ, Martinez S, Martin GR. 2008. Specific regions within the embryonic midbrain and cerebellum require different levels of FGF signaling during development. Development 135:889–898. doi:10.1242/DEV.011569

Blanco-Ameijeiras J, Lozano-Fernández P, Martı E. 2022. Centrosome maturation - in tune with the cell cycle. J Cell Sci 135. doi:10.1242/JCS.259395

Blom HJ, Shaw GM, Den Heijer M, Finnell RH. 2006. Neural tube defects and folate: case far from closed. Nat Rev Neurosci 7:724–731. doi:10.1038/NRN1986

Chambers SM, Fasano CA, Papapetrou EP, Tomishima M, Sadelain M, Studer L. 2009. Highly efficient neural conversion of human ES and iPS cells by dual inhibition of SMAD signaling. Nat Biotechnol 27:275–280. doi:10.1038/NBT.1529

Cheng C, Cong Q, Liu Y, Hu Y, Liang G, Dioneda KMM, Yang Y. 2023. Yap controls notochord formation and neural tube patterning by integrating mechanotransduction with FoxA2 and Shh expression. Sci Adv 9. doi:10.1126/SCIADV.ADF6927

Chenn A, Zhang YA, Chang BT, McConnell SK. 1998. Intrinsic polarity of mammalian neuroepithelial cells. Mol Cell Neurosci 11:183–193. doi:10.1006/mcne.1998.0680

Del Corral RD, Olivera-Martinez I, Goriely A, Gale E, Maden M, Storey K. 2003. Opposing FGF and retinoid pathways control ventral neural pattern, neuronal differentiation, and segmentation during body axis extension. Neuron 40:65–79. doi:10.1016/S0896-6273(03)00565-8

Deng Q, Andersson E, Hedlund E, Alekseenko Z, Coppola E, Panman L, Millonig JH, Brunet JF, Ericson J, Perlmann T. 2011. Specific and integrated roles of Lmx1a, Lmx1b and Phox2a in ventral midbrain development. Development 138:3399–3408. doi:10.1242/DEV.065482

de Vries M, Carpinelli M, Rutland E, Hatzipantelis A, Partridge D, Auden A, Anderson PJ, De Groef B, Wu H, Osterwalder M, Visel A, Jane SM, Dworkin S. 2020. Interrogating the Grainyhead-like 2 (Grhl2) genomic locus identifies an enhancer element that regulates palatogenesis in mouse. Dev Biol 459:194–203. doi:10.1016/j.ydbio.2019.11.015

Edri S, Hayward P, Jawaid W, Arias AM. 2019. Neuro-mesodermal progenitors (NMPs): a comparative study between pluripotent stem cells and embryo-derived populations. Development 146:dev180190. doi:10.1242/DEV.180190

Elosegui-Artola A, Andreu I, Beedle AEM, Lezamiz A, Uroz M, Kosmalska AJ, Oria R, Kechagia JZ, Rico-Lastres P, Le Roux AL, Shanahan CM, Trepat X, Navajas D, Garcia-Manyes S, Roca-Cusachs P. 2017. Force Triggers YAP Nuclear Entry by Regulating Transport across Nuclear Pores. Cell 171:1397–1410.e14. doi:10.1016/j.cell.2017.10.008

Eura N, Matsui TK, Luginbühl J, Matsubayashi M, Nanaura H, Shiota T, Kinugawa K, Iguchi N, Kiriyama T, Zheng C, Kouno T, Lan YJ, Kongpracha P, Wiriyasermkul P, Sakaguchi YM, Nagata R, Komeda T, Morikawa N, Kitayoshi F, Jong M, Kobashigawa S, Nakanishi M, Hasegawa M, Saito Y, Shiromizu T, Nishimura Y, Kasai T, Takeda M, Kobayashi H, Inagaki Y, Tanaka Y, Makinodan M, Kishimoto T, Kuniyasu H, Nagamori S, Muotri AR, Shin JW, Sugie K, Mori E. 2020. Brainstem Organoids From Human Pluripotent Stem Cells. Front Neurosci 14. doi:10.3389/FNINS.2020.00538

Gonzalez-Gobartt E, Blanco-Ameijeiras J, Usieto S, Allio G, Benazeraf B, Martí E. 2021. Cell intercalation driven by SMAD3 underlies secondary neural tube formation. Dev Cell 56:1147–1163.e6. doi:10.1016/j.devcel.2021.03.023

Gouti M, Delile J, Stamataki D, Wymeersch FJ, Huang Y, Kleinjung J, Wilson V, Briscoe J. 2017. A Gene Regulatory Network Balances Neural and Mesoderm Specification during Vertebrate Trunk Development. Dev Cell 41:243–261.e7. doi:10.1016/j.devcel.2017.04.002

Gouti M, Tsakiridis A, Wymeersch FJ, Huang Y, Kleinjung J, Wilson V, Briscoe J. 2014. In Vitro Generation of Neuromesodermal Progenitors Reveals Distinct Roles for Wnt Signalling in the Specification of Spinal Cord and Paraxial Mesoderm Identity. PLoS Biol 12:e1001937. doi:10.1371/JOURNAL.PBIO.1001937

Gribaudo S, Robert R, van Sambeek B, Mirdass C, Lyubimova A, Bouhali K, Ferent J, Morin X, van Oudenaarden A, Nedelec S. 2024. Self-organizing models of human trunk organogenesis recapitulate spinal cord and spine co-morphogenesis. Nat Biotechnol 42:1243–1253. doi:10.1038/S41587-023-01956-9

Haremaki T, Metzger JJ, Rito T, Ozair MZ, Etoc F, Brivanlou AH. 2019. Self-organizing neuruloids model developmental aspects of Huntington’s disease in the ectodermal compartment. Nat Biotechnol 37:1198–1208. doi:10.1038/S41587-019-0237-5;SUBJMETA

Ha YJJ, Nisal A, Tang I, Lee C, Jhamb I, Wallace C, Howarth R, Schroeder S, Vong KI, Meave N, Jiwani F, Barrows C, Lee S, Jiang N, Patel A, Bagga K, Banka N, Friedman L, Blanco FA, Yu S, Rhee S, Jeong HS, Plutzer I, Major MB, Benoit B, Poüs C, Heffner C, Kibar Z, Bot GM, Northrup H, Au KS, Strain M, Ashley-Koch AE, Finnell RH, Le JT, Meltzer HS, Araujo C, Machado HR, Stevenson RE, Yurrita A, Mumtaz S, Ahmed A, Khara MH, Mutchinick OM, Medina-Bereciartu JR, Hildebrandt F, Melikishvili G, Marwan AI, Capra V, Noureldeen MM, Salem AMS, Issa MY, Zaki MS, Xu L, Lee JE, Shin D, Alkelai A, Shuldiner AR, Kingsmore SF, Murray SA, Gee HY, Miller WT, Tolias KF, Wallingford JB, Tkemaladze T, Kirmani S, Gonda DD, Güneş AS, Kara B, Hanak B, Phillips HW, Salimi-Dafsari H, Takahashi Y, Shril S, Kolvenbach CM, Magana T, Araújo C, Lupo PJ, Au KS, Koch AEA, Kim S, Gleeson JG. 2025. The contribution of de novo coding mutations to meningomyelocele. Nature 641:419–426. doi:10.1038/S41586-025-08676-X

Hirata H, Tomita K, Bessho Y, Kageyama R. 2001. Hes1 and Hes3 regulate maintenance of the isthmic organizer and development of the mid/hindbrain. EMBO J 20:4454. doi:10.1093/EMBOJ/20.16.4454

Hsu PC, Yang CT, Jablons DM, You L. 2018. The Role of Yes-Associated Protein (YAP) in Regulating Programmed Death-Ligand 1 (PD-L1) in Thoracic Cancer. Biomedicines 6. doi:10.3390/BIOMEDICINES6040114

Iyer NR, Shin J, Cuskey S, Tian Y, Nicol NR, Doersch TE, Seipel F, McCalla SG, Roy S, Ashton RS. 2022. Modular derivation of diverse, regionally discrete human posterior CNS neurons enables discovery of transcriptomic patterns. Sci Adv 8. doi:10.1126/SCIADV.ABN7430

Jo J, Xiao Y, Sun AX, Cukuroglu E, Tran HD, Göke J, Tan ZY, Saw TY, Tan CP, Lokman H, Lee Y, Kim D, Ko HS, Kim SO, Park JH, Cho NJ, Hyde TM, Kleinman JE, Shin JH, Weinberger DR, Tan EK, Je HS, Ng HH. 2016. Midbrain-like Organoids from Human Pluripotent Stem Cells Contain Functional Dopaminergic and Neuromelanin-Producing Neurons. Cell Stem Cell 19:248–257. doi:10.1016/j.stem.2016.07.005

Kadoshima T, Sakaguchi H, Nakano T, Soen M, Ando S, Eiraku M, Sasai Y. 2013. Self-organization of axial polarity, inside-out layer pattern, and species-specific progenitor dynamics in human ES cell-derived neocortex. Proc Natl Acad Sci U S A 110:20284–20289. doi:10.1073/PNAS.1315710110

Karzbrun E, Khankhel AH, Megale HC, Glasauer SMK, Wyle Y, Britton G, Warmflash A, Kosik KS, Siggia ED, Shraiman BI, Streichan SJ. 2021. Human neural tube morphogenesis in vitro by geometric constraints. Nature 599:268–272. doi:10.1038/S41586-021-04026-9

Kasai T, Suga H, Sakakibara M, Ozone C, Matsumoto R, Kano M, Mitsumoto K, Ogawa K, Kodani Y, Nagasaki H, Inoshita N, Sugiyama M, Onoue T, Tsunekawa T, Ito Y, Takagi H, Hagiwara D, Iwama S, Goto M, Banno R, Takahashi J, Arima H. 2020. Hypothalamic Contribution to Pituitary Functions Is Recapitulated In Vitro Using 3D-Cultured Human iPS Cells. Cell Rep 30:18–24.e5. doi:10.1016/j.celrep.2019.12.009

Koch F, Scholze M, Wittler L, Schifferl D, Sudheer S, Grote P, Timmermann B, Macura K, Herrmann BG. 2017. Antagonistic Activities of Sox2 and Brachyury Control the Fate Choice of Neuro-Mesodermal Progenitors. Dev Cell 42:514–526.e7. doi:10.1016/j.devcel.2017.07.021

Krammer T, Stuart HT, Gromberg E, Ishihara K, Cislo D, Melchionda M, Becerril Perez F, Wang J, Costantini E, Lehr S, Arbanas L, Hörmann A, Neumüller RA, Elvassore N, Siggia E, Briscoe J, Kicheva A, Tanaka EM. 2024. Mouse neural tube organoids self-organize floorplate through BMP-mediated cluster competition. Dev Cell 59:1940–1953.e10. doi:10.1016/j.devcel.2024.04.021

Kumamoto T, Hanashima C. 2017. Evolutionary conservation and conversion of Foxg1 function in brain development. Dev Growth Differ 59:258–269. doi:10.1111/DGD.12367

Lancaster MA, Corsini NS, Wolfinger S, Gustafson EH, Phillips AW, Burkard TR, Otani T, Livesey FJ, Knoblich JA. 2017. Guided self-organization and cortical plate formation in human brain organoids. Nat Biotechnol 35:659–666. doi:10.1038/NBT.3906

Lancaster MA, Knoblich JA. 2014. Generation of cerebral organoids from human pluripotent stem cells. Nat Protoc 9:2329–2340. doi:10.1038/NPROT.2014.158

Lee JH, Shin H, Shaker MR, Kim HJ, Park SH, Kim JH, Lee N, Kang M, Cho S, Kwak TH, Kim JW, Song MR, Kwon SH, Han DW, Lee S, Choi SY, Rhyu IJ, Kim H, Geum D, Cho IJ, Sun W. 2022. Production of human spinal-cord organoids recapitulating neural-tube morphogenesis. Nat Biomed Eng 6:435–448. doi:10.1038/S41551-022-00868-4

Libby ARG, Joy DA, Elder NH, Bulger EA, Krakora MZ, Gaylord EA, Mendoza-Camacho F, Butts JC, McDevitt TC. 2021. Axial elongation of caudalized human organoids mimics aspects of neural tube development. Development 148. doi:10.1242/DEV.198275

Lindsay SJ, Xu Y, Lisgo SN, Harkin LF, Copp AJ, Gerrelli D, Clowry GJ, Talbot A, Keogh MJ, Coxhead J, Santibanez-Koref M, Chinnery PF. 2016. HDBR Expression: A Unique Resource for Global and Individual Gene Expression Studies during Early Human Brain Development. Front Neuroanat 10. doi:10.3389/FNANA.2016.00086

Marshall AR, Galea GL, Copp AJ, Greene NDE. 2023. The surface ectoderm exhibits spatially heterogenous tension that correlates with YAP localisation during spinal neural tube closure in mouse embryos. Cells and Development 174. doi:10.1016/j.cdev.2023.203840

Marthiens V, ffrench-Constant C. 2009. Adherens junction domains are split by asymmetric division of embryonic neural stem cells. EMBO Rep 10:515–520. doi:10.1038/EMBOR.2009.36

Maury Y, Côme J, Piskorowski RA, Salah-Mohellibi N, Chevaleyre V, Peschanski M, Martinat C, Nedelec S. 2015. Combinatorial analysis of developmental cues efficiently converts human pluripotent stem cells into multiple neuronal subtypes. Nat Biotechnol 33:89–96. doi:10.1038/NBT.3049

McMahon JA, Takada S, Zimmerman LB, Fan CM, Harland RM, McMahon AP. 1998. Noggin-mediated antagonism of BMP signaling is required for growth and patterning of the neural tube and somite. Genes Dev 12:1438–1452. doi:10.1101/GAD.12.10.1438

McNeely PD, Howes WJ. 2004. Ineffectiveness of dietary folic acid supplementation on the incidence of lipomyelomeningocele: pathogenetic implications. J Neurosurg 100:98–100. doi:10.3171/PED.2004.100.2.0098

Nakano T, Ando S, Takata N, Kawada M, Muguruma K, Sekiguchi K, Saito K, Yonemura S, Eiraku M, Sasai Y. 2012. Self-formation of optic cups and storable stratified neural retina from human ESCs. Cell Stem Cell 10:771–785. doi:10.1016/j.stem.2012.05.009

Nardone G, Oliver-De La Cruz J, Vrbsky J, Martini C, Pribyl J, Skládal P, Pešl M, Caluori G, Pagliari S, Martino F, Maceckova Z, Hajduch M, Sanz-Garcia A, Pugno NM, Stokin GB, Forte G. 2017. YAP regulates cell mechanics by controlling focal adhesion assembly. Nat Commun 8. doi:10.1038/NCOMMS15321

Ogura T, Sakaguchi H, Miyamoto S, Takahashi J. 2018. Three-dimensional induction of dorsal, intermediate and ventral spinal cord tissues from human pluripotent stem cells. Development 145. doi:10.1242/DEV.162214

Olivera-Martinez I, Harada H, Halley PA, Storey KG. 2012. Loss of FGF-dependent mesoderm identity and rise of endogenous retinoid signalling determine cessation of body axis elongation. PLoS Biol 10. doi:10.1371/JOURNAL.PBIO.1001415

O’Rahilly R, Müller F. 1987. Stages in Early Human Development. Future Aspects in Human In Vitro Fertilization 238–244. doi:10.1007/978-3-642-71412-2_33

Paridaen JTML, Wilsch-Bräuninger M, Huttner WB. 2013. XAsymmetric inheritance of centrosome-associated primary cilium membrane directs ciliogenesis after cell division. Cell 155:333. doi:10.1016/j.cell.2013.08.060

Pasca AM, Sloan SA, Clarke LE, Tian Y, Makinson CD, Huber N, Kim CH, Park JY, O’Rourke NA, Nguyen KD, Smith SJ, Huguenard JR, Geschwind DH, Barres BA, Pasca SP. 2015. Functional cortical neurons and astrocytes from human pluripotent stem cells in 3D culture. Nat Methods 12:671–678. doi:10.1038/NMETH.3415

Piccolo S, Dupont S, Cordenonsi M. 2014. The biology of YAP/TAZ: hippo signaling and beyond. Physiol Rev 94:1287–1312. doi:10.1152/PHYSREV.00005.2014

Porter FD, Drago J, Xu Y, Cheema SS, Wassif C, Huang SP, Lee E, Grinberg A, Massalas JS, Bodine D, Alt F, Westphal H. 1997. Lhx2, a LIM homeobox gene, is required for eye, forebrain, and definitive erythrocyte development. Development 124:2935–2944. doi:10.1242/DEV.124.15.2935

Rayon T, Maizels RJ, Barrington C, Briscoe J. 2021. Single-cell transcriptome profiling of the human developing spinal cord reveals a conserved genetic programme with human-specific features. Development 148. doi:10.1242/DEV.199711

Ribes V, Le Roux I, Rhinn M, Schuhbaur B, Dollé P. 2009. Early mouse caudal development relies on crosstalk between retinoic acid, Shh and Fgf signalling pathways. Development 136:665–676. doi:10.1242/DEV.016204

Rito T, Libby ARG, Demuth M, Domart MC, Cornwall-Scoones J, Briscoe J. 2025. Timely TGFβ signalling inhibition induces notochord. Nature 637:673–682. doi:10.1038/S41586-024-08332-W;TECHMETA

Saade M, Blanco-Ameijeiras J, Gonzalez-Gobartt E, Martı E. 2018. A centrosomal view of CNS growth. Development 145. doi:10.1242/DEV.170613

Saade M, Ferrero DS, Blanco-Ameijeiras J, Gonzalez-Gobartt E, Flores-Mendez M, Ruiz-Arroyo VM, Martínez-Sáez E, Ramón y Cajal S, Akizu N, Verdaguer N, Martí E. 2020. Multimerization of Zika Virus-NS5 Causes Ciliopathy and Forces Premature Neurogenesis. Cell Stem Cell 27:920–936.e8. doi:10.1016/j.stem.2020.10.002

Saade M, Gonzalez-Gobartt E, Escalona R, Usieto S, Martí E. 2017. Shh-mediated centrosomal recruitment of PKA promotes symmetric proliferative neuroepithelial cell division. Nat Cell Biol 19:493–503. doi:10.1038/NCB3512

Saade M, Martí E. 2025. Early spinal cord development: from neural tube formation to neurogenesis. Nat Rev Neurosci 26:195–213. doi:10.1038/S41583-025-00906-5

Saitsu H, Shiota K. 2008. Involvement of the axially condensed tail bud mesenchyme in normal and abnormal human posterior neural tube development. Congenit Anom (Kyoto) 48:1–6. doi:10.1111/J.1741-4520.2007.00178.X

Saitsu H, Yamada S, Uwabe C, Ishibashi M, Shiota K. 2004. Development of the posterior neural tube in human embryos. Anat Embryol (Berl) 209:107–117. doi:10.1007/S00429-004-0421-2

Santos C, Murray A, Marshall AR, Metcalfe K, Narayan P, Castro SCP de, Maniou E, Greene NDE, Galea GL, Copp AJ. 2023. Spinal neural tube formation and regression in human embryos. Elife 12. doi:10.7554/ELIFE.88584.1

Simunovic M, Metzger JJ, Etoc F, Yoney A, Ruzo A, Martyn I, Croft G, You DS, Brivanlou AH, Siggia ED. 2019. A 3D model of a human epiblast reveals BMP4-driven symmetry breaking. Nat Cell Biol 21:900–910. doi:10.1038/S41556-019-0349-7

Takemoto T, Uchikawa M, Kamachi Y, Kondoh H. 2006. Convergence of Wnt and FGF signals in the genesis of posterior neural plate through activation of the Sox2 enhancer N-1. Development 133:297–306. doi:10.1242/DEV.02196

Tang C, Wang X, D’Urso M, van der Putten C, Kurniawan NA. 2022. 3D Interfacial and Spatiotemporal Regulation of Human Neuroepithelial Organoids. Adv Sci (Weinh) 9. doi:10.1002/ADVS.202201106

Turner DA, Girgin M, Alonso-Crisostomo L, Trivedi V, Baillie-Johnson P, Glodowski CR, Hayward PC, Collignon J, Gustavsen C, Serup P, Steventon B, Lutolf MP, Arias AM. 2017. Anteroposterior polarity and elongation in the absence of extra-embryonic tissues and of spatially localised signalling in gastruloids: mammalian embryonic organoids. Development 144:3894–3906. doi:10.1242/DEV.150391

Tzouanacou E, Wegener A, Wymeersch FJ, Wilson V, Nicolas JF. 2009. Redefining the Progression of Lineage Segregations during Mammalian Embryogenesis by Clonal Analysis. Dev Cell 17:365–376. doi:10.1016/j.devcel.2009.08.002

Varelas X, Sakuma R, Samavarchi-Tehrani P, Peerani R, Rao BM, Dembowy J, Yaffe MB, Zandstra PW, Wrana JL. 2008. TAZ controls Smad nucleocytoplasmic shuttling and regulates human embryonic stem-cell self-renewal. Nat Cell Biol 10:837–848. doi:10.1038/NCB1748

Varelas X, Samavarchi-Tehrani P, Narimatsu M, Weiss A, Cockburn K, Larsen BG, Rossant J, Wrana JL. 2010. The Crumbs Complex Couples Cell Density Sensing to Hippo-Dependent Control of the TGF-β-SMAD Pathway. Dev Cell 19:831–844. doi:10.1016/j.devcel.2010.11.012

Verrier L, Davidson L, Gierlinśki M, Dady A, Storey KG. 2018. Neural differentiation, selection and transcriptomic profiling of human neuromesodermal progenitor-like cells in vitro. Development 145. doi:10.1242/DEV.166215

Vong KI, Lee S, Au KS, Crowley TB, Capra V, Martino J, Haller M, Araújo C, Machado HR, George R, Gerding B, James KN, Stanley V, Jiang N, Alu K, Meave N, Nidhiry AS, Jiwani F, Tang I, Nisal A, Jhamb I, Patel Arzoo, Patel Aakash, McEvoy-Venneri J, Barrows C, Shen C, Ha YJ, Howarth R, Strain M, Ashley-Koch AE, Azam M, Mumtaz S, Bot GM, Finnell RH, Kibar Z, Marwan AI, Melikishvili G, Meltzer HS, Mutchinick OM, Stevenson DA, Mroczkowski HJ, Ostrander B, Schindewolf E, Moldenhauer J, Zackai EH, Emanuel BS, Garcia-Minaur S, Nowakowska BA, Stevenson RE, Northrup H, McNamara HK, Aldinger KA, Phelps IG, Deng M, Glass IA, Morrow B, McDonald-McGinn DM, Sanna-Cherchi S, Lamb DJ, Gleeson JG, Meltzer HS, Au KS, Magana T, Yurrita A, Zaki MS, Mumtaz S, Medina-Bereciartu JR, Kolvenbach CM, Shril S, Hildebrandt F, Noureldeen MM, Salem AMS, Takahashi Y, Salimi-Dafsari H, Phillips HW, Hanak B, Kara B, Güneş AS, Gonda DD, Kirmani S, Tkemaladze T. 2024. Risk of meningomyelocele mediated by the common 22q11.2 deletion. Science 384:584–590. doi:10.1126/SCIENCE.ADL1624

Wilson PA, Hemmati-Brivanlou A. 1995. Induction of epidermis and inhibition of neural fate by Bmp-4. Nature 376:331–333. doi:10.1038/376331A0

Wilson V, Olivera-Martinez I, Storey KG. 2009. Stem cells, signals and vertebrate body axis extension. Development 136:1591–1604. doi:10.1242/DEV.021246

Wolujewicz P, Aguiar-Pulido V, AbdelAleem A, Nair V, Thareja G, Suhre K, Shaw GM, Finnell RH, Elemento O, Ross ME. 2021. Genome-wide investigation identifies a rare copy-number variant burden associated with human spina bifida. Genetics in Medicine 23:1211–1218. doi:10.1038/s41436-021-01126-9

Xue X, Kim YS, Ponce-Arias AI, O’Laughlin R, Yan RZ, Kobayashi N, Tshuva RY, Tsai YH, Sun S, Zheng Y, Liu Y, Wong FCK, Surani A, Spence JR, Song H, Ming GL, Reiner O, Fu J. 2024. A patterned human neural tube model using microfluidic gradients. Nature 628:391–399. doi:10.1038/S41586-024-07204-7

Yan CH, Levesque M, Claxton S, Johnson RL, Ang SL. 2011. Lmx1a and lmx1b function cooperatively to regulate proliferation, specification, and differentiation of midbrain dopaminergic progenitors. J Neurosci 31:12413–12425. doi:10.1523/JNEUROSCI.1077-11.2011

Zeng B, Liu Z, Lu Y, Zhong S, Qin S, Huang L, Zeng Y, Li Z, Dong H, Shi Y, Yang J, Dai Y, Ma Q, Sun L, Bian L, Han D, Chen Y, Qiu X, Wang W, Marín O, Wu Q, Wang Y, Wang X. 2023. The single-cell and spatial transcriptional landscape of human gastrulation and early brain development. Cell Stem Cell 30:851–866.e7. doi:10.1016/j.stem.2023.04.016

Zheng Y, Xue X, Resto-Irizarry AM, Li Z, Shao Y, Zheng Y, Zhao G, Fu J. 2019. Dorsal-ventral patterned neural cyst from human pluripotent stem cells in a neurogenic niche. Sci Adv 5:5933–5944. doi:10.1126/SCIADV.AAX5933

