## Supplementary Figures for "HUMAN SPINAL CORD ORGANOIDS REVEAL CELL INTERCALATION AS A CONSERVED MECHANISM FOR SECONDARY NEURULATION"

Blanco-Ameijeiras, Supplementary Figure 1

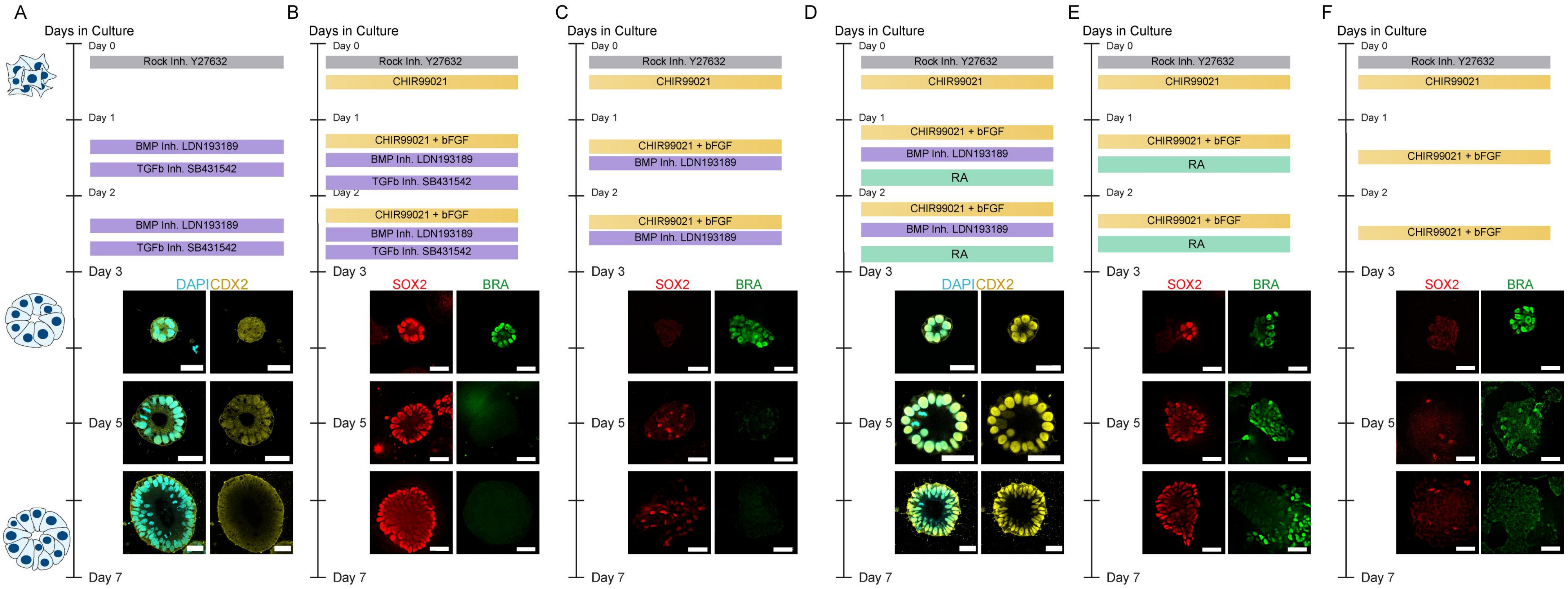

Blanco-Ameijeiras, Supplementary Figure 2

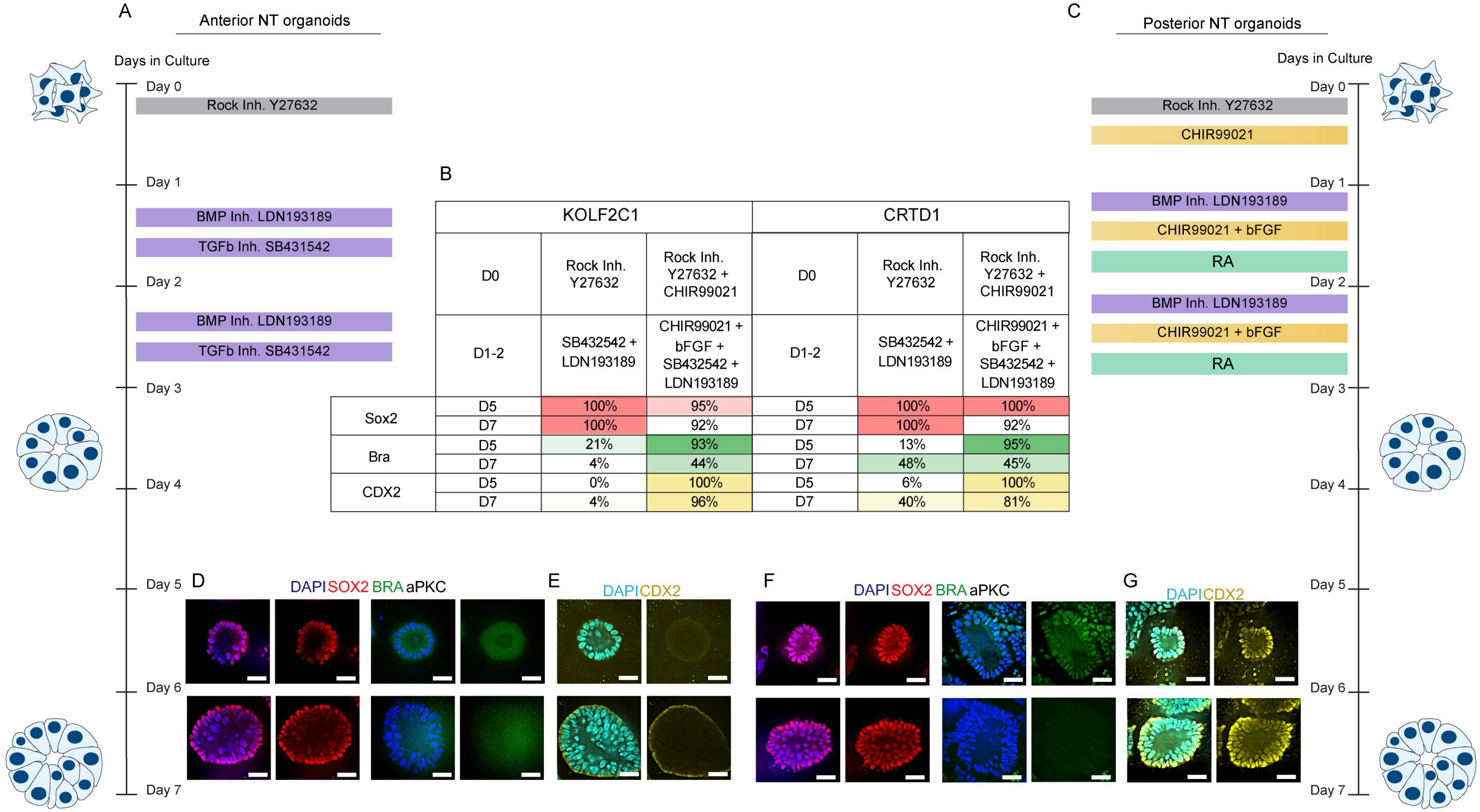


| | | KOLF2C1 | | | CRTD1 | |
|---|---|---|---|---|---|---|
| | D0 | Rock Inh. Y27632 | Rock Inh. Y27632 + CHIR99021 | D0 | Rock Inh. Y27632 | Rock Inh. Y27632 + CHIR99021 |
| | D1-2 | SB432542 + LDN193189 | CHIR99021 + bFGF + SB432542 + LDN193189 | D1-2 | SB432542 + LDN193189 | CHIR99021 + bFGF + SB432542 + LDN193189 |
| Sox2 | D5 | 100% | 95% | D5 | 100% | 100% |
| | D7 | 100% | 92% | D7 | 100% | 92% |
| Bra | D5 | 21% | 93% | D5 | 13% | 95% |
| | D7 | 4% | 44% | D7 | 48% | 45% |
| CDX2 | D5 | 0% | 100% | D5 | 6% | 100% |
| | D7 | 4% | 96% | D7 | 40% | 81% |

Blanco-Ameijeiras, Supplementary Figure 3

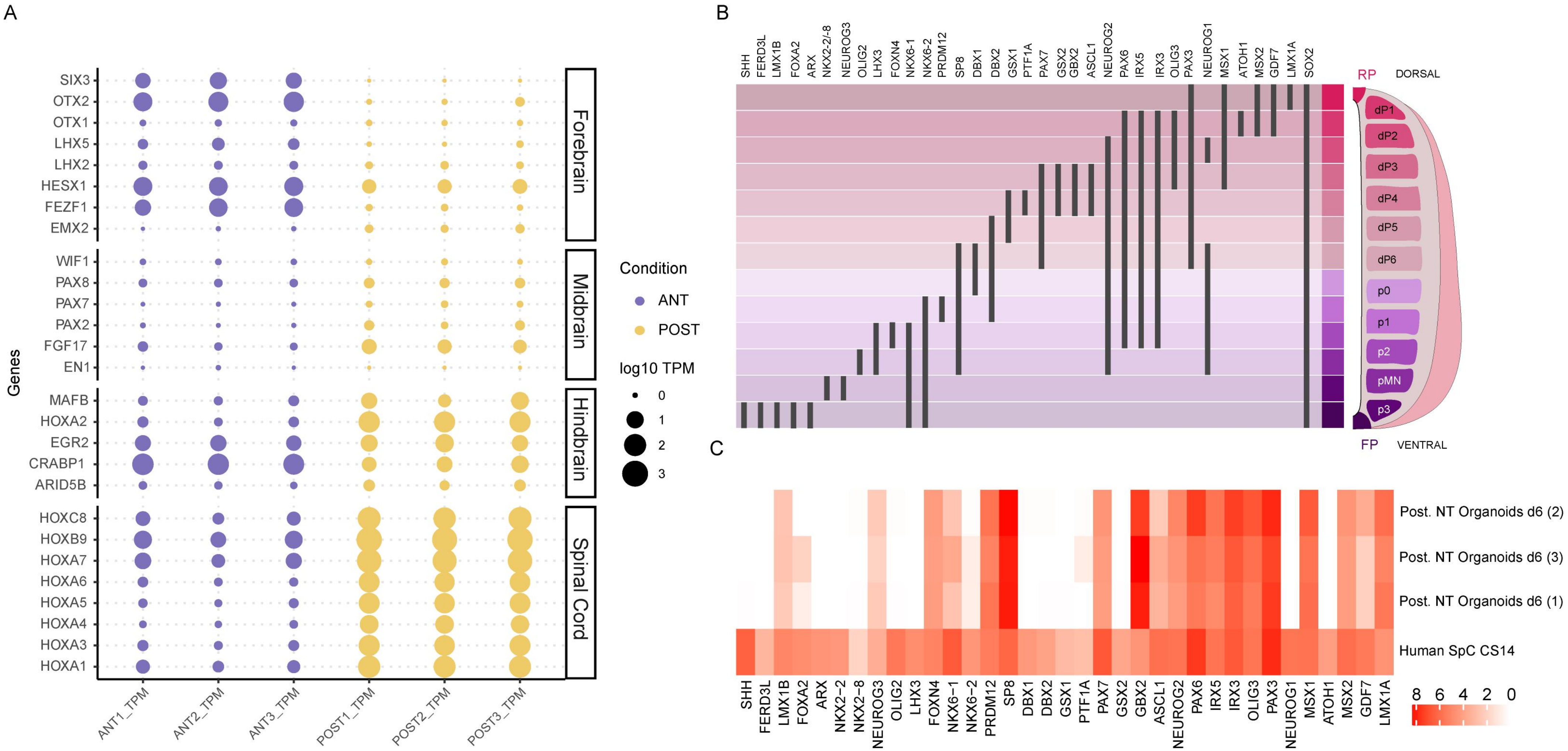

Blanco-Ameijeiras, Supplementary Figure 4

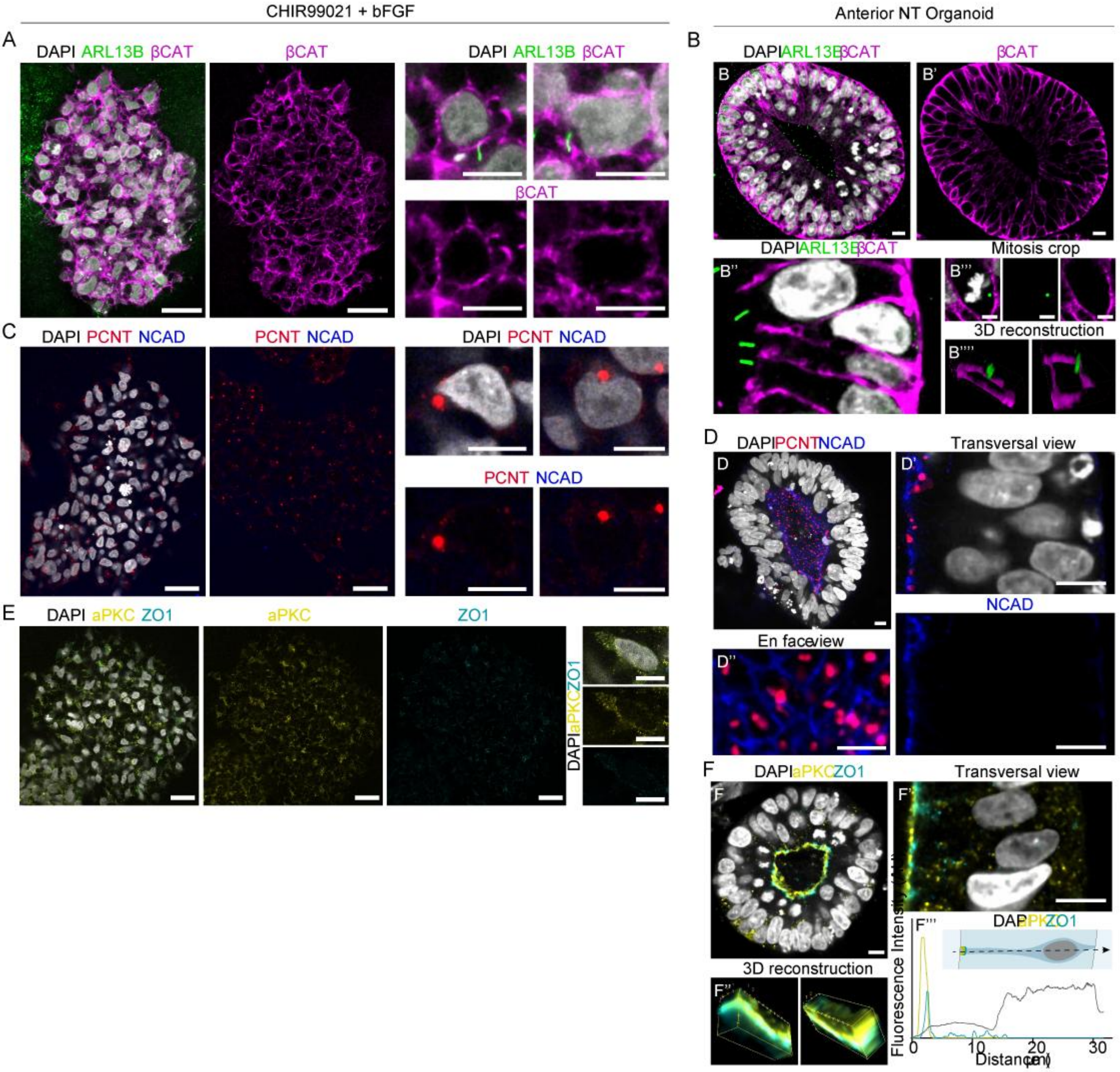

Blanco-Ameijeiras, Supplementary Figure 5

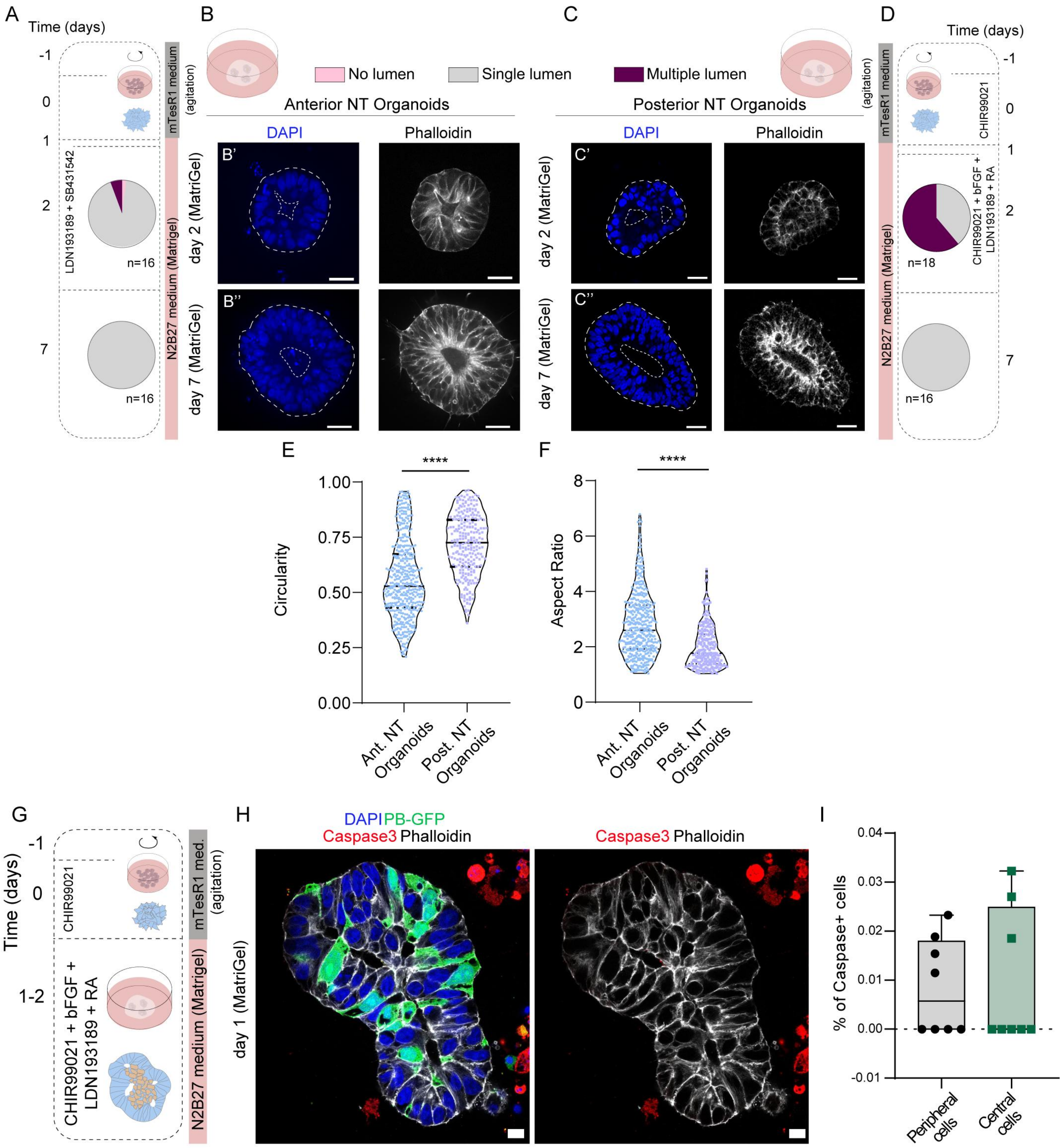

### Blanco-Ameijeiras, Supplementary Figure 6

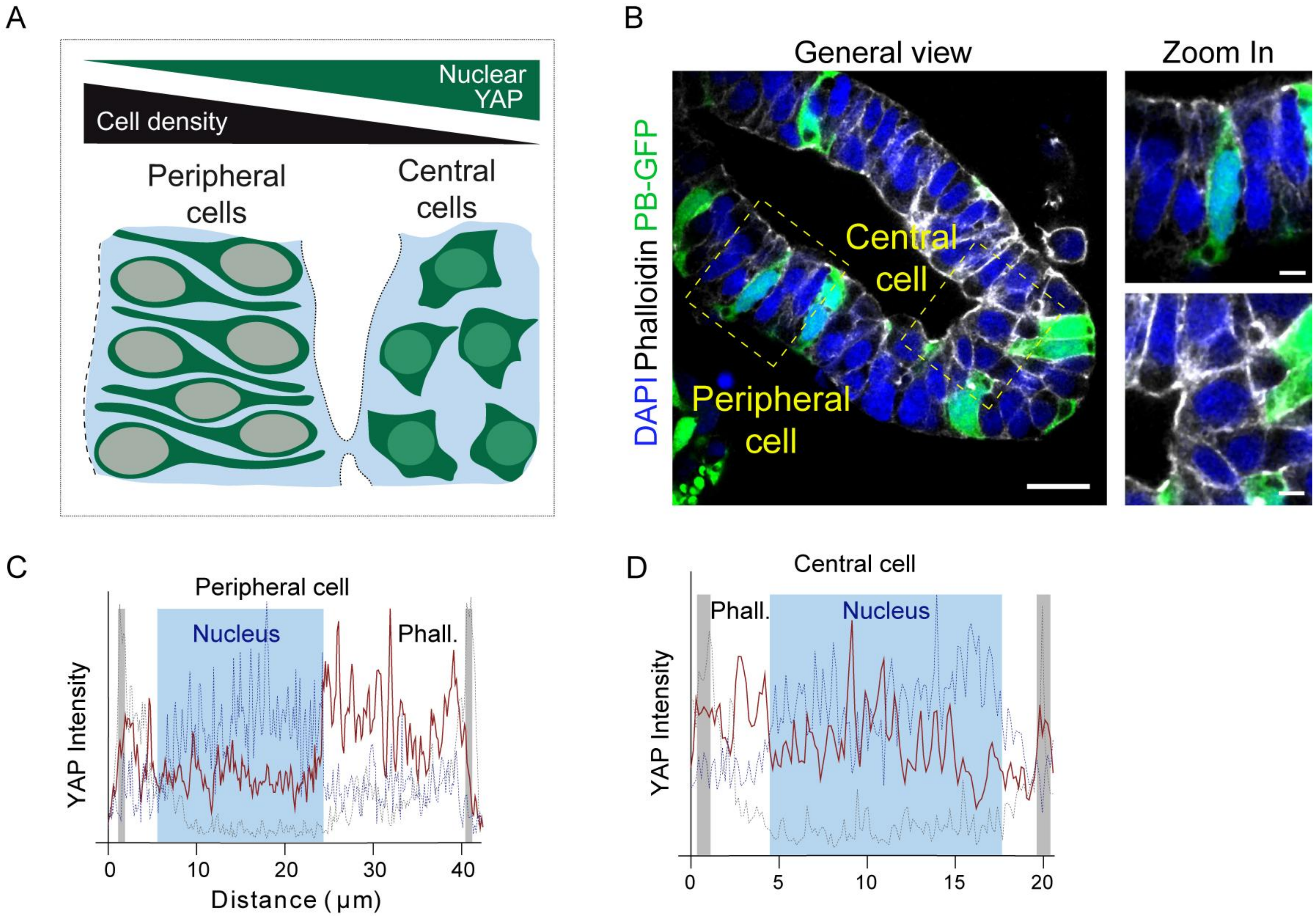
